# Cost-effective DNA Storage System with DNA Movable Type

**DOI:** 10.1101/2024.07.17.603163

**Authors:** Chenyang Wang, Di Wei, Zheng Wei, Dongxin Yang, Jing Xing, Yunze Wang, Xiaotong Wang, Pei Wang, Guannan Ma, Xinru Zhang, Haolan Li, Chuan Tang, Pengfei Hou, Jie Wang, Renjun Gao, Guiqiu Xie, Cuidan Li, Yingjiao Ju, Peihan Wang, Liya Yue, Yongliang Zhao, Yongjie Sheng, Jingfa Xiao, Haitao Niu, Sihong Xu, Huaiyi Yang, Di Liu, Bo Duan, Dongbo Bu, Guangming Tan, Fei Chen

**Author notes:** Correspondence to (Fei Chen), (Guangming Tan), (Dongbo Bu), (Bo Duan). These authors contributed equally to this work (Chenyang Wang, Di Wei, Zheng Wei, Dongxin Yang, Jing Xing, Yunze Wang, Xiaotong Wang).

## Abstract

In the face of exponential data growth, DNA-based storage offers a promising solution for preserving big-data. However, most existing DNA storage methods, akin to traditional block printing, require costly chemical synthesis for each individual data file, adopting a sequential, one-time-use synthesis approach. To overcome these limitations, we introduce a novel, cost-effective "DNA-Movable-Type Storage" system, inspired by movable type printing. This system utilizes pre-fabricated DNA movable types-short, double-stranded DNA oligonucleotides encoding specific payload, address, and checksum data. These DNA-MTs are enzymatically ligated/assembled into cohesive sequences, termed "DNA movable type blocks", streamlining the assembly process with the automated BISHENG-1 DNA-MT inkjet printer. Using BISHENG-1, we successfully printed, assembled, stored and accurately retrieved 43.7 KB of data files in diverse formats (text, image, audio, and video) *in vitro* and *in vivo*, using only 350 DNA-MTs. Notably, each DNA-MT, synthesized once (2 *OD*), can be used up to 10,000 times, reducing costs to 122 $/MB-outperforming existing DNA storage methods. This innovation circumvents the need to synthesize entire DNA sequences encoding files from scratch, offering significant cost and efficiency advantages. Furthermore, it has considerable untapped potential to advance a robust DNA storage system, better meeting the extensive data storage demands of the big-data era.

## 1 Introduction

In the era of big data, diverse data, such as text, image, audio, and video, are being generated at an unprecedented rate, with predictions that global data will exceed 380 zettabytes by 2028. [1] However, current storage media (hard disk, magnetic tape, *etc*) are unable to meet the storage demands due to their large physical size, limited lifetime (up to around 30 years), and high energy consumption, resulting in an estimation that less than 2% of this data will be preserved. [2] Consequently, new techniques suitable for large-size data storage are highly desired.

DNA-based data storage presents a promising solution to big data storage, offering high density, longevity, and low energy consumption. [3] For instance, the human genome, containing over 3 giga base pairs, weighs only 3 picograms; [4] moreover, successful sequencing ancient genomes like the mammoth’s, [5] underscores DNA’s potential for preserving data over millennia with minimal energy. These remarkable attributes make DNA-based storage an ideal candidate for addressing large-scale and long-term data storage challenges.

Since 2010, advancements in DNA-based data storage technology have rapidly accelerated, driven by breakthroughs in DNA sequencing and synthesis technologies. [6] Key milestones include the introduction of “bit-to-base” encoding by Church’s group in 2012 and Goldman’s group in 2013, which facilitates the translation of digital data into DNA sequences. [7] Since then, various sophisticated encoding schemes have emerged, incorporating error correction capabilities and strategies to overcome biological constraints. Notable examples include the fountain code, which optimizes data redundancy and approaches Shannon capacity while providing robustness against corruption; [8] the yin-yang codec system, which balances robustness, compatibility, and information density for DNA data storage; [9] the composite DNA letters that enable DNA-based data storage using fewer DNA synthesis cycles; [10] and LDPC codes combined with pseudo-random sequences in a superposition coding process have also enhanced scalability by allowing artificial chromosomes to tolerate high-error reads from MinION sequencing. [11] Additionally, innovations like DNA-Aeon, [12] Explorer, [13] and Wukong [14] provide customizable encoding interfaces, allowing users to control GC content, homopolymer run length, and undesired motifs, while maintaining high coding efficiency. Notably, the establishment of the “DNA Data Storage Alliance” in 2020 represents a significant step towards commercial viability, aiming to unify and propel industry efforts. [15]

Despite these advances, it remains experimental due to the high DNA synthesis cost, substantially constraining its practicality. [16] Extensive researches are ongoing to reduce costs by improving synthesis efficiency through column- or microarray-based methods and exploring enzyme-mediated synthesis. [17] Although these improvements have significantly lowered the costs of DNA storage, the current expenses associated with DNA-based storage remain too high for practical application.

A closer look reveals that most existing DNA storage methods resemble woodblock printing, necessitating costly and time-consuming chemical synthesis for each data file from scratch. This sequential (base-by-base), one-time-use synthesis approach significantly hinders its practicality, highlighting an urgent need for more efficient and reusable strategies.

Addressing the bottlenecks, our study draws inspiration from ancient Chinese movable type printing to innovatively propose “DNA movable type (DNA-MT) storage”. Unlike traditional one-time-use approaches prevalent in existing DNA storage technologies, this method employs reusable, pre-fabricated “DNA movable types (DNA-MTs)”, which significantly lowers DNA synthesis and storage costs. Additionally, the enzymatic ligation process can be easily parallelized, further increasing its efficiency and scalability.

Our exploration into DNA movable type storage began in 2017. The first-generation DNA movable type comprised a middle movable type sequence (12 nt for payload, 8 nt for address), flanked by conserved PCR primer sequences (15 nt) and two restriction enzyme sites (*Bam*HⅠ and *Sbf*Ⅰ) (**Figure S1**). Five DNA-MTs were sequentially linked to assembly a DNA-MT block through one or multiple PCR steps, which comprised one payload type and four address types. Both ends of the DNA-MT block was equipped with two restriction enzyme sites to facilitate cloning into pUC19 plasmids for *in vitro* and *in vivo* storage (**Figure S1b**). Using our first-generation DNA-MT design, we successfully stored the Tang poem "The Road Is Difficult" (232 bytes).[18]

In recent two years, other research teams have also made innovative contributions to DNA movable type (DNA-MT) storage. Yuan *et al.* developed the DNA-MT structure by reducing the DNA-MT sequence to 6 nucleotides and extending the PCR primer arms to 43 nucleotides. [19] They also incorporated two Type IIS restriction sites at both ends to facilitate subsequent ligation. Liu *et al.* proposed a novel technique that utilizes polymerase-catalyzed primer exchange reactions, where specific primers are extended from hairpin DNA templates. The extended primers then undergo branch migration, allowing multiple DNA-MT blocks to be connected together. [20] In 2024, epigenetic modifications were explored as a new method for data storage. The ’epi-bit’ system uses DNA-MT sequences with methylation sites on CpG dinucleotides within single-stranded DNA, where cytosine can be either methylated or unmethylated. DNMT1 enzyme recognizes partially methylated sites and adds a methyl group to the complementary cytosine, encoding an epi-bit as ’1’. [21]

The above attempts demonstrated the feasibility of DNA-MT storage scheme, but also highlighted limitations, such as complex operations and low storage density. To address these issues, we began transitioning to a second-generation design in 2020, moving away from the PCR-based movable type block assembly approach used in the first generation.

This manuscript presents our work on a second-generation DNA-MT storage solution, offering a promising approach for practical DNA data storage. This method combines one-step enzymatic ligation with DNA inkjet printing to assemble DNA-MT blocks. We start by using pre-made double-stranded oligonucleotides, called “DNA-MTs,” which are designed to encode specific payloads or address data. Here, we also introduced column- and row-checksum DNA-MTs for error detection and correction. Following this, using a DNA-MT code table, we selected the appropriate pre-made DNA-MTs for payloads, addresses, and checksums of target files, and enzymatically ligated them into cohesive sequences termed “DNA-MT blocks”. To automate the block assembly process, we engineered a DNA-MT inkjet printer, “BISHENG-1”, which handles the selection, injection, and ligation of DNA-MTs. [22] Following assembly, these blocks are directly cloned into plasmids for preservation, either *in vitro* or *in vivo* in *E. coli*.

Utilizing our second-generation DNA-MT storage system, we successfully achieved DNA-MT printing, storage (both *in vitro* and *in vivo*), and 100% accurate decoding of 43.7 KB of text, images, audio, and video data (**Table S1**) using only 350 DNA-MTs. This technology, in conjunction with the "BISHENG-1" system, demonstrate cost advantages in data writing, enabling the DNA-MTs to be used/printed up to 10,000 times after a single synthesis (2 *OD*). Currently, with a throughput of 556 and a printing speed of 4 bytes/second, the cost can be reduced to 122 $/MB, surpasses all reported DNA storage technologies to the best of our knowledge. Additionally, BISHENG-1 printed 43.7 KB in ∼5 hours, showcasing its efficiency advantage over sequential one-time-use synthesis methods (∼100 hours). Looking ahead, this technology holds significant untapped potential for further cost and efficiency improvements, promising to evolve a more robust DNA storage ecosystem to meet the rapidly growing storage demands of the big data era.

## 2 Results

### 2.1 DNA Movable Type storage principle

Drawing inspiration from ancient Chinese movable type printing, DNA-MT storage employs a comparable technique for data encoding and storage, exemplified by *Shakespeare’s Sonnet 12* (**Figure 1**).

**Figure 1.**
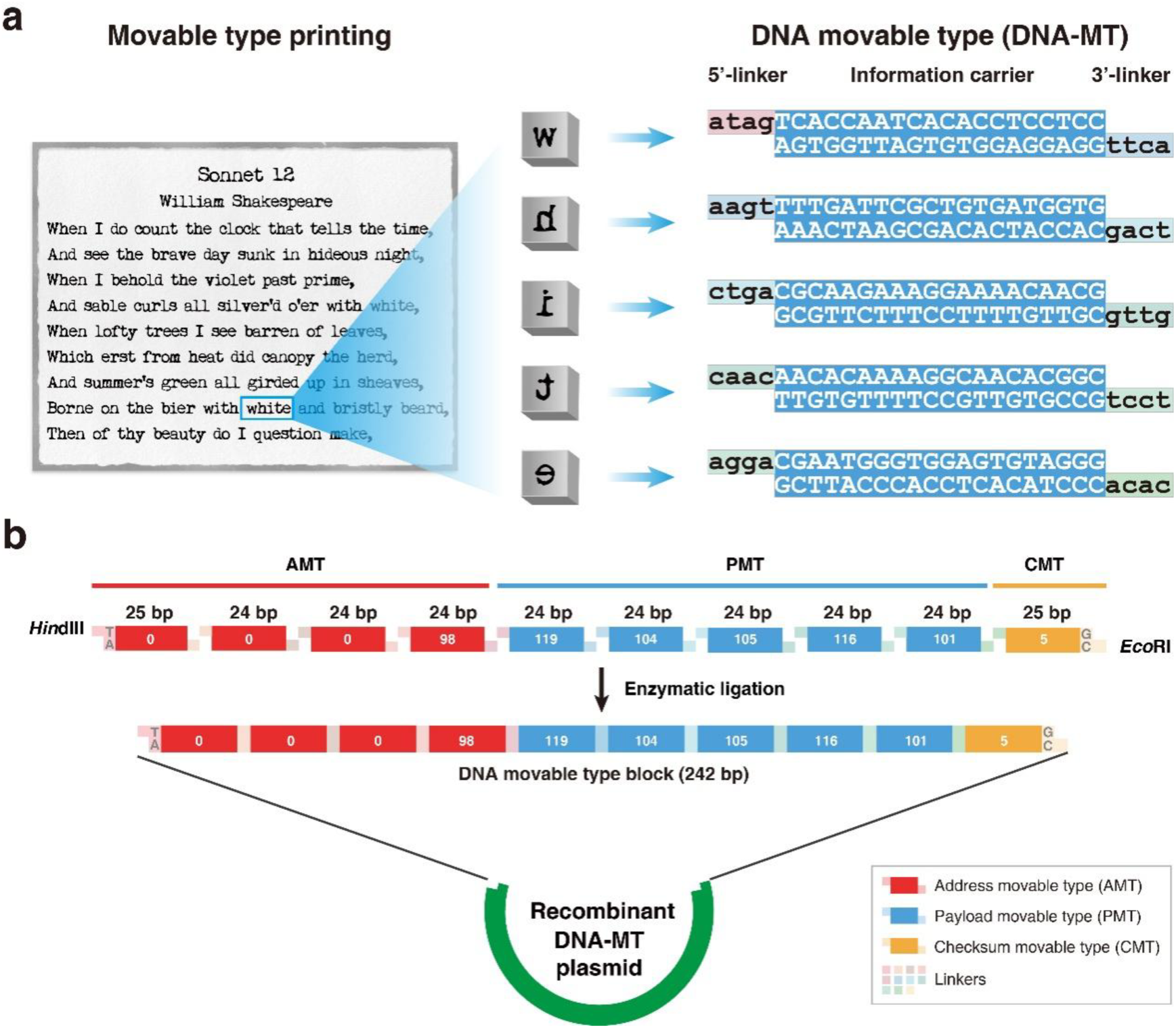
DNA-MT storage principle and structure. **a**, Mapping relationship between movable type printing and DNA-MT storage exemplified by the word “white” from *Shakespeare’s Sonnet 12*. The process of assembling pre-made wooden types in a frame to print text (left panel) is compared to the assembly of pre-synthesized DNA- MT, where each MT corresponds to a character (right panel). The word “white” from *Shakespeare’s Sonnet 12* is used to illustrate the method. Each character in the word "white" is represented by a DNA-MT consisting of a 20-bp double-stranded DNA information carrier with two 4-nt sticky ends that facilitate sequential assembly to form in a cohesive DNA sequence that encodes “white”. **b**, Structure of DNA-MT block designed to encode the word "white". This block contains five Payload DNA-MTs (PMTs) encoding ASCII codes for “white” (119, 104, 105, 116, 101), four Address DNA-MTs (AMTs) specifying a four-byte address (0, 0, 0, 98) indicative of the 340^th^ character position, and a checksum DNA-MT (CMT) for error detection (set at 5). These ten DNA-MTs are enzymatically sequentially linked using T4 DNA ligase through 11 linkers to form a cohesive DNA-MT block, with *Eco*RI and *Hin*dIII linkers at the ends for cloning into DNA-MT plasmid, enabling both *in vitro* and *in vivo* storage.

Historically, the replacement of woodblock printing by movable type printing greatly facilitated the spread of culture and the advancement of civilization. This was mainly due to its significant advantages in cost and efficiency over woodblock printing. Movable type printing is based on reusable pre-fabricated wooden or metal types for each character, allowing for their free selection and assembly into a frame to print texts like *Sonnet 12* (**Figure 1a**, left panel). This approach enables the efficient assembly of pages using pre-made characters, significantly reducing the costs and labor required for printing. In contrast, woodblock printing is a single-use, costly, and labor-intensive method that requires a unique carved block for each page. This method lacks the reusability and flexibility of movable type printing, making it more expensive and time-consuming.

Drawing on the evolution from woodblock to movable type printing, we have innovated an analogous method in the realm of DNA-based storage, termed DNA movable type storage (**Figure 1**). DNA-MTs, serving as the cornerstone of this method, are pre-synthesized, short, double-stranded DNA fragments (20-nt, termed “information carriers”), each flanked by sticky ends (4-nt, termed “linkers”) to facilitate easy linkage. Each DNA-MT encodes a byte of payload/address/checksum information. The method enables the selective ligation of multiple DNA-MTs into a cohesive, extended sequence, forming a “DNA-MT block” with a distinct address and checksum, thus improving the DNA storage’s precision. These blocks are then directly cloned into the optimized *pUC19* plasmid (termed “DNA-MT-plasmid”) (details in Methods section) using *Eco*RI and *Hin*dIII restriction sites on the outermost DNA-MTs, enabling preservation either *in vitro* within the plasmid or *in vivo* through *E. coli* transformation. Similar to the reusability and flexibility of movable type printing, the DNA-MT storage method offers significant cost and efficiency advantages.

**Figure 1a** illustrates five payload DNA-MTs (PMTs) within a DNA-MT block, encoding the word “white” from *Sonnet 12*. The first PMT, encoding “w”, contains an “information carrier” sequence “TCACC…CCTCC/AGTGG…GGAGG”, with its 3’- end linker “*ttca*” pairing with the 5’-end linker “*aagt*” of the second PMT for “h”. Similarly, the 3’-end linker of the second type pairs with the 5’-end linker of the third one (for “i”), continuing this pattern to yield a DNA sequence “*atag*TCACCAATCACACCTCCTCC*aagt*…CGAATGGGTGGAGTGTAGGG” that represents “white”. In addition to these PMTs, four address DNA-MTs (AMTs) and a checksum DNA-MT (CMT) are integrated for precise localization and error correction (**Figure 1b**). Employing T4 DNA ligase, these ten DNA-MTs are ligated via nine linkers into a cohesive double-stranded oligonucleotide block (DNA-MT block) in a single step, encoding the word “white” at the 340^th^ character address (notated as “0, 0, 0, 98”) with a checksum of 5 at the end. Notably, the two block’s terminal linkers, *Eco*RI and *Hin*dIII restriction sites, facilitate direct cloning the block into the DNA-MT-plasmid for *in vitro* storage or transformation into *E. coli* for *in vivo* storage. Moreover, our DNA-MT storage technology exhibits good versatility and scalability, as evidenced by our successful encoding and storage of a diverse range of file formats beyond text, including images, audio, and video (detailed in **TableS1**).

### 2.2 DNA-MT inkjet printer for automating block assembly: BISHENG-1

To automate DNA-MT storage process, we developed BISHENG-1, a DNA-MT inkjet printer capable of constructing DNA-MT blocks with high-throughput. BISHENG-1 consists of five main components (**Figure 2**): multiple cartridges for holding DNA- MTs, pre-digested vectors (*Hin*dIII and *Eco*RI) and T4 ligase, a conveyor belt for tube transport, an automated tube loading and discharging device, a printing monitor and a programable control system. The printer is currently equipped with 556 cartridges that contain varying amounts of DNA-MTs (AMTs, PMTs, and CMTs) according to specific requirements, along with the digested vector and ligase. Each tube on the conveyor is designed to collect ten DNA-MTs of a block plus the digested vector and ligase for subsequent enzymatic ligation into a long oligonucleotide.

**Figure 2.**
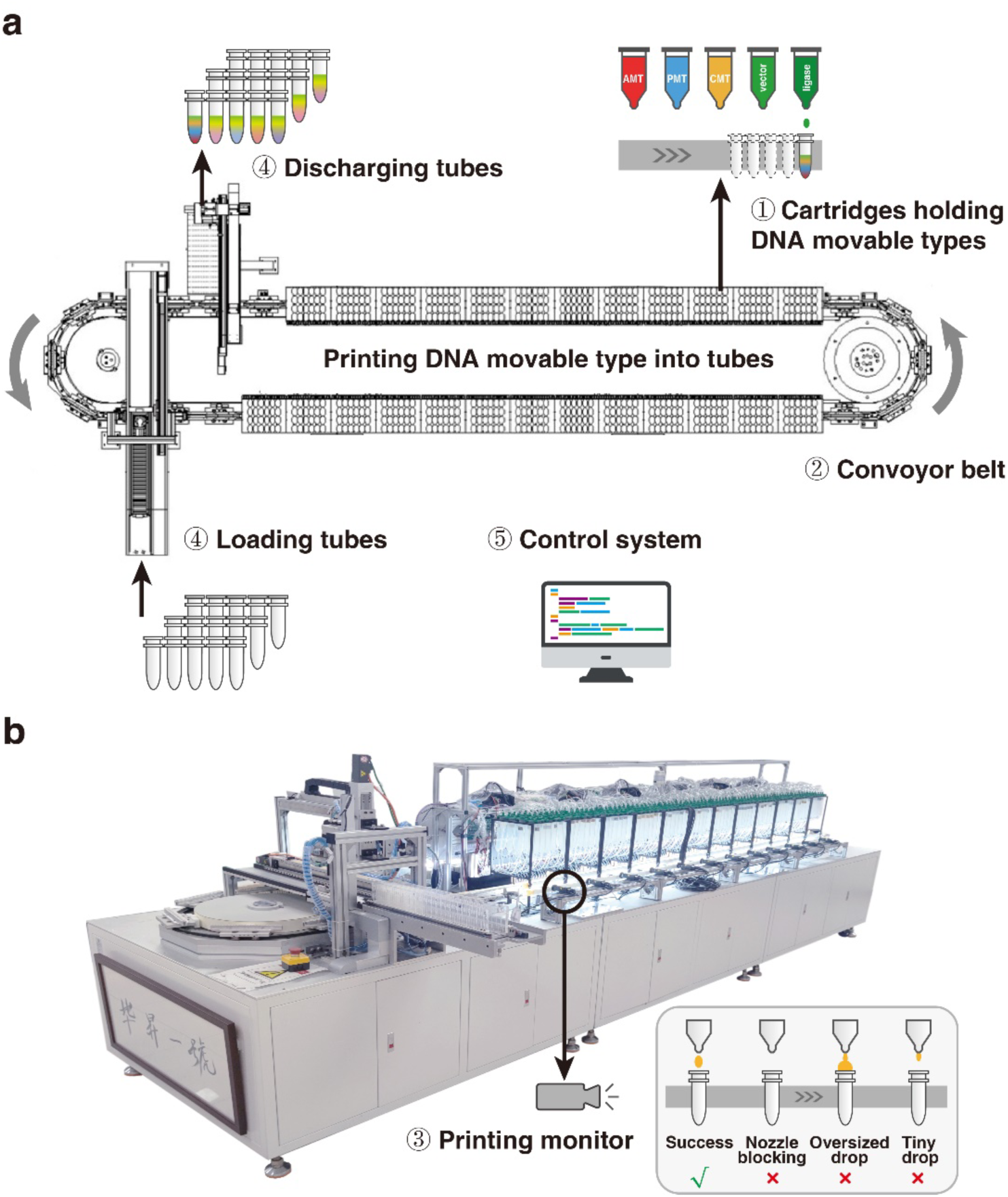
DNA movable type inkjet printer BISHENG-1 for automated DNA-MT block assembly. a, Architecture of BISHENG-1. The printer includes: i) multiple cartridges holding DNA movable types, pre-digested vectors, and T4 ligase, essential for DNA-MT assembly; ii) a conveyor belt for carrying and transporting tubes; iii) a printing monitor for real-time feedback and error detection; iv) devices for automatic tube loading and unloading; v) a programmable control system. Each tube on the conveyor is designed to collect all DNA-MTs of a block plus the digested vector and ligase for subsequent one-step enzymatic ligation into a cohesive DNA-MT block. **b,** A photograph of BISHENG-1 with a focus on the printing monitor.

BISHENG-1 operates like standard inkjet printers, executing a cycle of “*loading empty tubes, printing DNA-MTs into tubes, and discharging the filled tubes*” (**Figure 2a**; see **Supplementary Video**). Initially, empty tubes are placed onto the conveyor, moving past various cartridges. As tubes pass the cartridges for designated DNA-MTs, pre-digested vector and ligase, BISHSENG-1 commands the corresponding cartridge to inject a drop (∼6 µl) of each into the tubes. Once the tube has passed all cartridges, it has collected all necessary components: assigned DNA-MTs, digested vector, and ligase. Upon collection completion, the filled tubes are discharged, making room for a new cycle (loading, printing, and discharging), thereby streamlining the DNA storage process.

However, issues like cartridge nozzle blockages or incorrect droplet sizes may occur. To address these errors, BISHENG-1 is equipped with a video camera and image analysis software for monitoring malfunctioning nozzles (**Figure 2b**). Detected errors trigger automatic reprinting of affected tubes, ensuring a high success rate.

### 2.3 Workflow of BISHENG-1 DNA-MT storage

BISHENG-1 DNA-MT storage system comprises four key steps: encoding, assembling/printing, storage, and decoding, as detailed in **Figure 3**.

**Figure 3.**
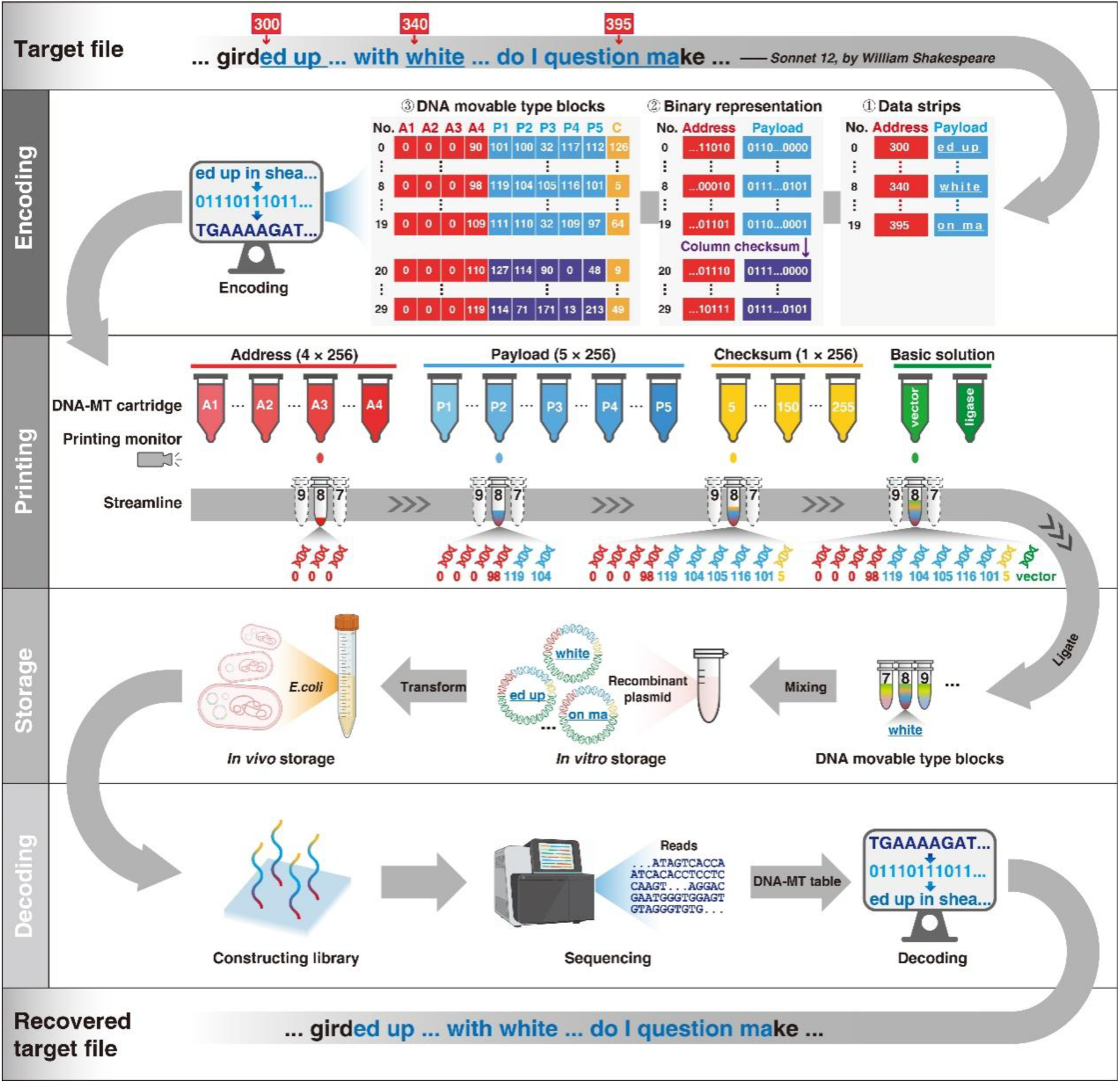
Workflow of DNA-MT storage. The main steps include: **(1) Encoding**: The target file, specifically *Sonnet 12*, is divided into 100-byte/character fragments. Each fragment is subdivided into 20 data strips, and each strip comprises a five-byte payload and a four-byte address. The figure illustrates 20 sequential data strips (rows 0 – 19) of the fourth stripe, covering payloads from the 300^th^ to the 399^th^ character (“ed up…question ma”), with the 8^th^ strip encoding “white” at the 340^th^ position. Additional error detection and correction are provided through column (rows 20 – 29) and row checksums (column 9). **(2) Printing**: Utilizing BISHENG-1 DNA-MT inkjet printer, ten specific DNA-MTs (4 AMTs, 5 PMTs, and 1 CMT), along with ligase and pre- digested vector are printed into each tube for enzymatic ligation into a DNA-MT block within plasmids. **(3) Storage**: The assembled recombinant plasmids are preserved either in liquid/lyophilized forms for *in vitro* storage or transformed into *E. coli* for long-term *in vivo* storage. **(4) Decoding**: Through next-generation sequencing, DNA- MT blocks are sequenced. The resulting DNA sequences are then decoded back into movable types according to the code table, thereby reconstructing the original data (e.g., decoding a 250 nt sequence enables the recovery of the word “white” at position 340). The decoded PMTs are arranged by their AMTs to restore the original file.

#### 2.3.1 Encoding the target file into DNA-MTs (Encoding)

##### Fragmentation, stripe formation, and binary conversion

First, we divide the target file (*Sonnet 12*) into 100-byte/character fragments, subsequently splitting each fragment into 20 addressed data strips to form a stripe. Each strip contains five bytes/characters (payload) and the address of the first character. **Figure 3** displays 20 consecutive data strips (row 0 – 19) of the fourth stripe, spanning payloads from the 300^th^ to 399^th^ character (“ed up…with white…do I question ma”). The first data strip encodes “ed up”, beginning at the 300^th^ character, and the last strip encodes “on ma”, starting at the 395^th^ character. The word “white”, introduced in the previous section, is in the 8^th^ data strip, beginning at the 340^th^ character position. The binary representations of these payload and address data (**Figure S2b**) are ultimately transformed into DNA-MT IDs presented in decimal format.

##### Error detection and correction

To address sequencing errors like base insertions, deletions and substitutions, and incorrect connections from enzymatic ligation failures, we introduce additional strips and DNA-MTs for error detection and correction. Utilizing the Reed-Solomon algorithm, [23] ten column-checksum strips (row 20 – 29) are generated from the previously mentioned twenty data strips (row 0 – 19) to identify and rectify column errors. Further, a row-checksum DNA-MT for each DNA-MT block (highlighted in orange) is calculated using the XOR algorithm, [24] enabling row-wise error detection (**Figure 3**, “Encoding” panel).

##### Translation to DNA-MT blocks

Employing a DNA-MT code table, we transform each byte from data strips (including payload, address and checksum) into its corresponding DNA-MT, further forming DNA-MT blocks. In the example in **Figure 3**, each block comprises four AMTs (labeled as *A*_1_, *A*_2_, *A*_3_, *A*_4_), five PMTs (labeled as *P*_1_, *P*_2_, …, *P*_5_), and one CMT. Notably, each AMT (8 bits) represents one of 256 different values, therefore four AMTs in one block allow for the assignment of a unique address to each of the 256^4^ possible addresses. This configuration yields a storage capacity of up to 20 gigabytes (256^4^ strips × 5 bytes/strip). Certainly, the system’s storage capacity can be expanded with more AMTs and PMTs per block. Detailed encoding procedures are provided in the Methods section.

#### 2.3.2 Assembling DNA-MT blocks through enzymatic ligation (Assembling)

To construct a DNA-MT block, we enzymatically connect the ten DNA-MTs into a cohesive long oligonucleotide by joining the 11 linkers of the DNA-MTs together using T4 ligase. Taking the 8^th^ strip from the fourth stripe as an example, we linked four AMTs (*A*_1_-0, *A*_2_-0, *A*_3_-0, *A*_4_-98), five PMTs (*P*_1_-119, *P*_2_-104, *P*_3_-105, *P*_4_-116, *P*_5_-101), and one CMT (*C* -5), yielding a cohesive 242 bp oligonucleotide sequence “…atagTCACCAATCACACCTCCTCCaagt…aggaCGAATGGGTGGAGTGTAGG Gtgtg…” (Detailed across **Figure 1 and 3**’s “Encoding panel” and fully in Methods section).

To automate assembling process, we developed a specialized DNA-MT inkjet printer equipped with 556 cartridges. BISHENG-1 inkjet printer operates similarly to a traditional inkjet printer but employs cartridges filled with DNA-MTs rather than ink, injecting these types into EP tubes instead of onto paper. Each EP tube is designated to collect all DNA-MTs for a specific block, e.g., the 8^th^ tube for the 8^th^ DNA-MT block’s ten DNA-MTs (**Figure 3**, “Printing” panel). As tubes pass cartridges with matching DNA-MTs, they’re automatically selected and filled. The process, including tube movement and the DNA-MT selection and injection, is fully automated. Two basic solutions (pre-digested vector and T4 DNA ligase) are then automatically injected into these tubes for enzymatic ligation, ultimately cloning the DNA-MTs into an DNA-MT- plasmid. A demonstrative video (**Supplementary Video**) offers an in-depth view of BISHENG-1 inkjet printer’s operation.

#### 2.3.3 *In vitro* and *in vivo* storage of DNA-MT blocks (Storage)

The recombinant plasmids from the above process are storable in liquid or lyophilized form for *in vitro* storage. These plasmids can be further introduced into *E. coli* for *in vivo* storage (**Figure 3**, “Storage” panel). *E. coli*’s rapid growth attribute ensures quick, reliable duplication of DNA-MT blocks in the recombinant plasmids through generations, providing a cost-effective and stable method for data replication and storage.

#### 2.3.4 Decoding DNA-MT stored file with error correction (Decoding)

To retrieve original files encoded within the plasmids, we use high-throughput sequencing to read the DNA-MT blocks. The sequences are then decoded corresponding movable types based on the DNA-MT code table. For example, a 250 nt DNA read “…atagTCACCAATCACACCTCCTCCaagt…aggaCGAATGGGTG…” is decoded into ten DNA-MTs. This includes four AMTs (*A*_1_-0, *A*_2_-0, *A*_3_-0, *A*_4_-98), five PMTs (*P*_1_-119, *P*_2_-104, *P*_3_-105, *P*_4_ -116, *P*_5_-101) and one CMT (*C* -5), reconstructing the original data strip “white” at starting address 340 (**Figure 3**, “Decoding” panel). The decoded blocks are then organized by address to recover the original file. Errors are detected and corrected using column-checksum strips and row- checksum DNA-MTs (details in Methods section).

### 2.4 DNA-MT design and library construction

The DNA-MT comprises an information carrier flanked by linkers at both ends. Using specific screening criteria (**Figure 4**), we optimized the sequences of both the linkers and the complete DNA-MT. This optimization is crucial for enhancing the efficiency and accuracy of DNA-MT ligation and assembly throughout the entire process.

**Figure 4.**
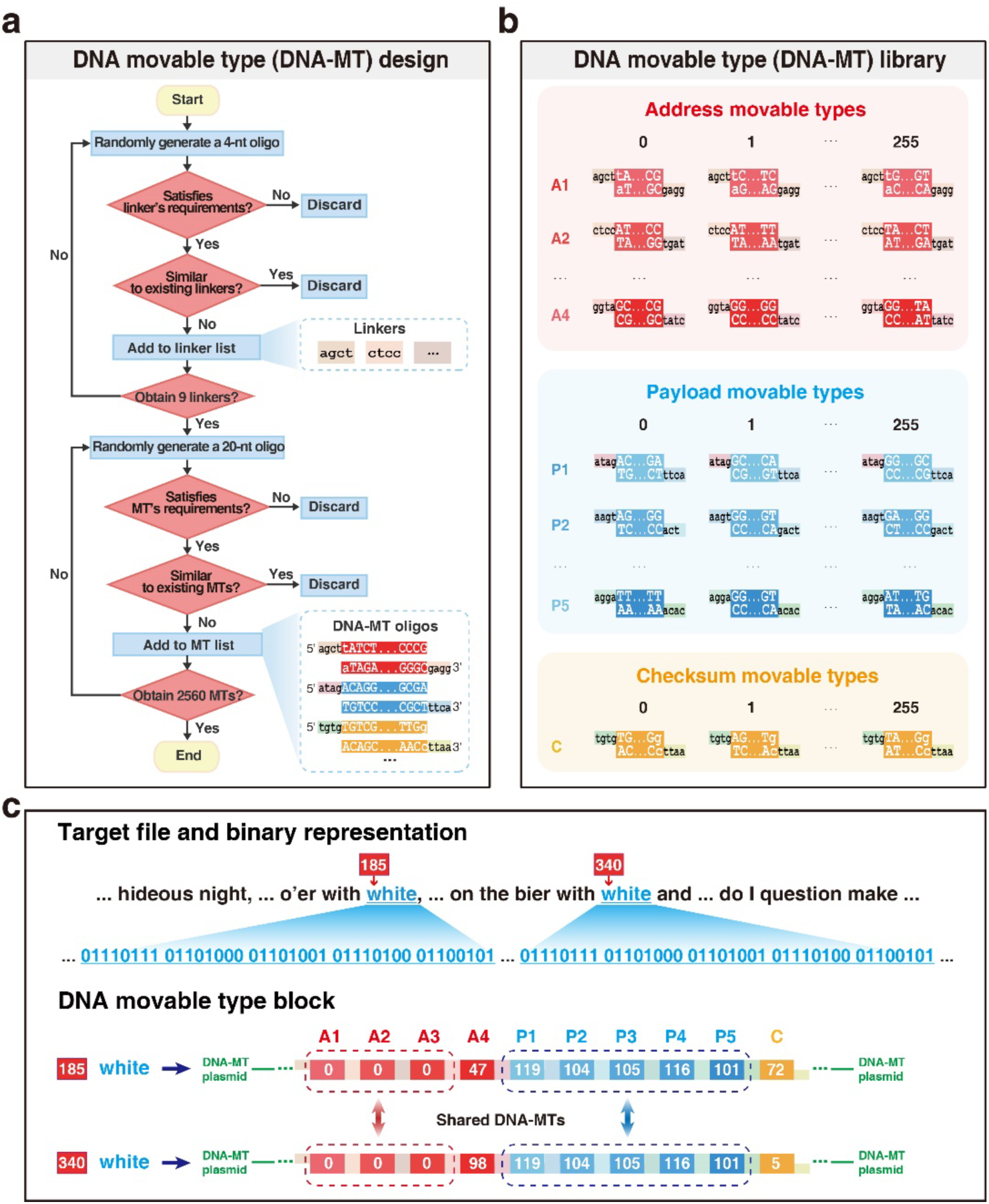
DNA-MT design and library construction. a, Sequence selection for DNA- MTs. 4-nt and 20-nt random oligonucleotides, as candidate linkers and information carriers for DNA-MTs, are subjected to a series of filtering criteria (such as GC content and homopolymer runs) to select optimal sequences, ensuring DNA-MTs’ functionality and specificity; **b,** DNA-MT Library Composition. The DNA-MT library is divided into three sub-libraries: AMT sub-library (1024 AMTs: 4×256), PMT sub-library (1280 PMTs: 5×256), and CMT sub-library (256 CMTs); **c,** Example of Pre-Made DNA-MTs for Reusability: Illustrates two DNA-MT blocks encoding the word “white” at positions 185 and 340. These blocks share three AMTs and five PMTs, exemplifying the reusability of pre-made DNA-MTs for reducing costs by minimizing the need for synthesizing new oligonucleotides for each occurrence.

#### 2.4.1 Linker design and assessment

We began by selecting linkers with high specificity to ensure precise pairings with their complementary partners. Non-specific linkers can lead to misassembly among DNA- MTs, such as the incorrect linkage of *A*_!_ with *A*_#_, instead of the correct *A*_"_, resulting in disorganized DNA-MT blocks. To maximize specificity, we used two main strategies. First, we applied strict criteria on sequence properties, including base diversity, homopolymer runs, and GC content, to reduce synthesis and sequencing errors. Linker lengths were also optimized, and 4-base linkers were selected for their balance between affinity and specificity. From a comprehensive screening of 256 possible 4-nt combinations (4^4^), we identified 200 candidate 4-base linkers (details in **Supplementary Table 2**) that met our stringent requirements for high specificity and accurate assembly. For cloning into pUC19 plasmids, we used *HindIII* and *EcoRI* restriction sites as fixed linkers at block ends (*agct, aatt*).

Using our established criteria, we developed a scoring system to evaluate linker effectiveness, filtering out those prone to misligation from the 200 candidates (see Methods for scoring details). Each DNA-MT block in our scheme contains 10 DNA- MTs connected by 11 linkers, with fixed restriction enzyme sites (*HindIII* and *EcoRI*) at both ends. Therefore, our scoring focused on selecting the nine internal linkers. We conducted 5,000 simulations to identify the optimal combinations of these nine linkers, achieving several combinations with specificity scores above 50, theoretically indicating a low likelihood of misassemblies (**Figure S3**). Validation experiments ultimately confirmed the currently used set of nine linkers with a specificity score of 52. We also compared this linker set with an alternative set scoring 55 in specificity. Both sets demonstrated nearly 100% block assembly accuracy (**Table S2**, **Table S3**, **Figure S4**).

#### 2.4.2 Information carrier design and assessment

Previous studies have shown that **high GC content, long homopolymer runs, and complex secondary structures** can increase synthesis and sequencing errors, while short editing distances among similar sequences elevate the risk of misdecoding [25]. **To optimize the functionality and specificity of the information carriers with linkers, we applied strict selection criteria** on GC content, homopolymer runs, secondary structures, and sequence editing distance [26]. To evaluate whether carrier length affects assembly accuracy, we conducted an experiment using eight blocks, each containing four DNA-MTs, with information carriers of 20 and 30 bp (DNA-MT sequences detailed in **Tables S4** and **S5**). As shown in **Figure S5, t**he 20-base DNA-MTs exhibited a higher exact match rate during assembly, suggesting improved precision with this length. Based on these findings, we selected 20-base carriers for subsequent experiments.

#### 2.4.3 Assembly experiment of 4 to 12 DNA-MT

To identify the optimal number of DNA-MTs for block assembly, we conducted preliminary experiments using blocks with 4 to 12 DNA-MTs (**Figure S6**). We observed that as the number of DNA-MTs increased, ligation/assembly accuracy slightly declined. Even with 12 DNA-MTs, the ligation accuracy of DNA-MT block assembly remains above 97%. To minimize potential errors associated with read length limitations, we prioritized configurations of full covered of all DNA-MTs by single sequencing reads. Thus, we selected a 10 DNA-MT scheme for subsequent experiments. This setup allows each 10 DNA-MT block (242 bp) to be fully covered by a single 250 nt read on the Illumina sequencing platform, eliminating the need for additional read assembly.

#### 2.4.4 DNA-MT library design and construction

After careful selection of the 11 4-nt linkers, we integrated these predetermined linkers with selected 20-nt information carrier candidates to complete the design of our DNA- MTs, based on the established criteria (detailed in the Methods section). Each DNA-MT in our scheme includes two linkers at the 3’ and 5’ ends, and a central information carrier encoding 1 byte of data, designated to encode data for payload, address, and checksum information.

Following the design of individual DNA-MTs, we proceeded to the design and assembly of DNA-MT blocks, each comprising 10 DNA-MTs, strategically configured to enhance data storage and retrieval efficiency. This configuration includes 5 PMTs encoding 5 bytes of data, 4 AMTs encoding 4 bytes of address information to support a total DNA-MT storage capacity of ∼2 GB, and each CMT encoding 1 byte of checksum for error correction. The blocks are constructed by linking the 5’ and 3’ ends of these DNA-MTs To ensure sequential ligation of DNA-MTs in a specified order *A*_1_, *A*_2_, *A*_3_, *A*_4_, *P*_1_, *P*_2_, *P*_3_, *P*_4_, *P*_5_, *C*, two strategic constraints were implemented (**Figure 4b**): i) All *A*_!_ DNA-MTs share identical linkers, and so do DNA-MTs in *A*_"_ to *C*; ii) The 3’-linker of an *A*_!_ DNA-MTs are designed to specifically pair with the 5’-linker of an *A*_"_ DNA-MTs, and so on.

Following these design principles, we successfully designed 2,560 unique DNA- MTs with high specificity, including 1,024 AMTs (4×256), 1,280 PMTs (5×256), and 256 CMTs. The detailed sequences of these carriers are systematically cataloged in a DNA-MT table (**Supplementary Table 1**) for ease access.

Employing pre-fabricated DNA-MTs significantly reduces synthesis from scratch. For instance, the word “white” occurs twice in *Sonnet 12* at addresses 185 and 340 (**Figure 4c**). Traditional “base-by-base” synthesis method would require repeating oligonucleotide synthesis for such case. However, with DNA-MT blocks, three AMTs and five PMTs are shared across recurring occurrences.

### 2.5 Data recovery robustness of the BISHENG-1 DNA-MT storage system

BISHENG-1 successfully printed and stored 43.7 KB of data (16,830 DNA-MT blocks) across four file formats (text, JPEG, MP3, and MP4) in approximately five hours (4 bytes/s) using only 350 unique DNA-MTs. In addition to *in vitro* storage, these recombinant plasmids containing DNA-MT blocks were also transformed into *E. coli* for *in vivo* storage, demonstrating data preservation within living cells.

To reconstruct the original files from DNA samples stored *in vivo* or *in vitro*, we sequenced the DNA samples and decoded the acquired sequences back into DNA-MT blocks. In each DNA-MT block, PMTs provide five bytes of data, while AMTs encode the address of these bytes. By compiling data decoded from all blocks, we successfully reconstructed the original files (Figure 5).

**Figure 5.**
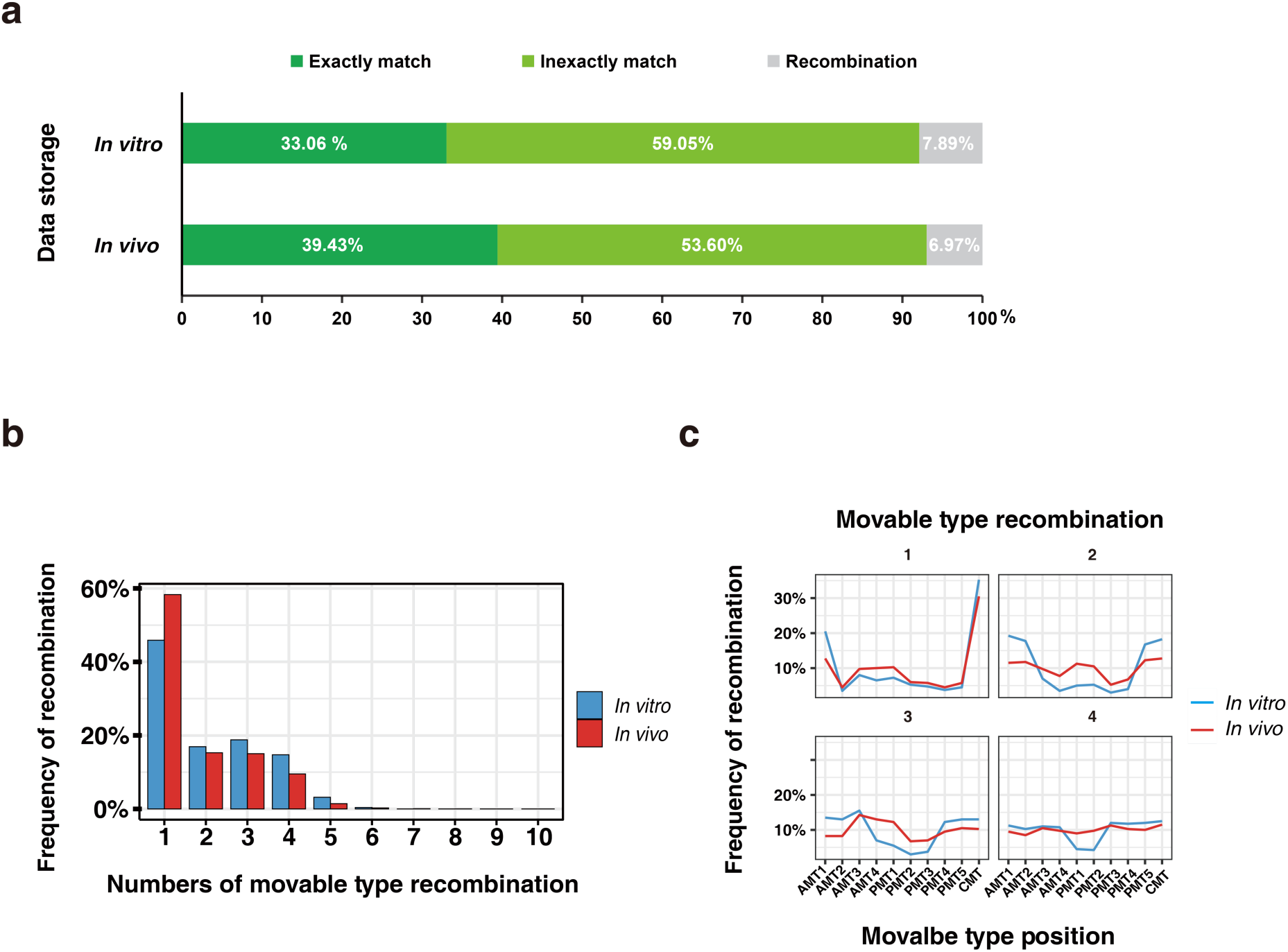
Data Recovery Robustness Analysis of the BISHENG-1 DNA-MT Storage System. **a**, Sequencing alignment results for files stored *in vivo* and *in vitro* using DNA-MTs. Exact matches indicate DNA-MTs with no mismatches, while inexactly matches allow for up to two base mismatches. Recombination refers to incorrect assembly events within a DNA-MT block. **b,** Proportion of recombination events within a DNA-MT block (composed of 10 DNA-MTs) under both *in vivo* and *in vitro* conditions. **c,** Position of recombination events within a DNA-MT block (composed of 10 DNA-MTs) under both *in vivo* and *in vitro* conditions.

During file reconstruction, errors could arise in the DNA-MT blocks, including missing DNA-MTs due to ligation failures, recombination during PCR, and missing blocks from insufficient sequencing coverage. We implemented row and column- checksum verification on the decoded DNA-MT blocks to address these issues (see Methods for details).

For *in vitro* storage, over 92% of sequencing reads were decoded correctly, recovering 16,794 out of the 16,830 DNA-MT blocks encoding the four target files. Of these, 33.06% were exact matches, and 59.05% were inexactly matches, while approximately 7.89% of reads contained errors due to block recombination. For *in vivo* storage, over 93% of reads were perfectly decoded, recovering 16,788 blocks, with 39.43% exact matches and 53.60% inexactly matches. Around 6.97% of reads had recombination errors, which were predominantly located at the ends of DNA blocks (**Figure 5c**). In both cases, any remaining DNA-MT blocks were recovered using the row-checksum algorithm, enabling the reconstruction of all four files with 100% accuracy.

### 2.6 Cost analysis of DNA-MT storage

The cost of storing data using DNA-MTs is quantified by the formula:

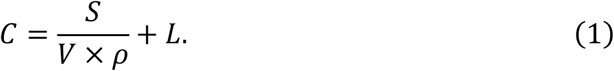

Here, *C* ($/byte) represents the cost per byte of data storage; *S* ($) denotes the synthesis cost of all DNA-MTs; *V* (μl) represents the total volume of DNA-MTs solution; 𝜌 (byte/μl) represents the information density (how many bytes of data can be encoded per μl of solution); 𝐿 ($/byte) represents the cost of T4 DNA ligase per byte. Based on BISHENG-1’s data, we define these parameters as follows:

#### Synthesis cost 𝑺 and solution volume 𝑽

*S* is 122.20 $ for synthesizing 470 OD of 5’-phosphorylated DNA-MTs at a commercial rate (∼260 $ for 1000 *OD*, *General Biol* (Anhui) Co. Ltd). The solution’s total volume is 2.4 ×10^7^ μl, setting *V* = 2.4 × 10^&^ μl.

#### Information density 𝝆

Assembling a DNA-MT block requires 60 μl solution (6 μl/block), and 30 blocks encode 80 bytes, yielding 𝜌 = 80/(60 × 30) = 0.044 bytes/μl.

#### Ligation cost 𝐿

Utilizing in-house produced T4 DNA ligase at approximately 0.43/g, the cost to ligate 1 MB is about 0.23 $ based on enzyme activity estimation. [27] Thus, 𝐿 = 2.19 × 10^’&^ $/byte.

Applying these parameter, the DNA-MT storage cost is estimated at *C* =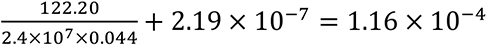 $/byte (122 $/MB), significantly lower than existing DNA storage technologies to the best of our knowledge (**Figure 6a**). The cost benefits of DNA-MTs become more pronounced for larger datasets, with costs diminishing to 35.28 $/MB for 1 GB and 30.56 $/MB for 1 TB (detailed in **Table S6**), thereby demonstrating the cost efficiency of DNA-MT storage through reusability and enzymatic ligation.

**Figure 6.**
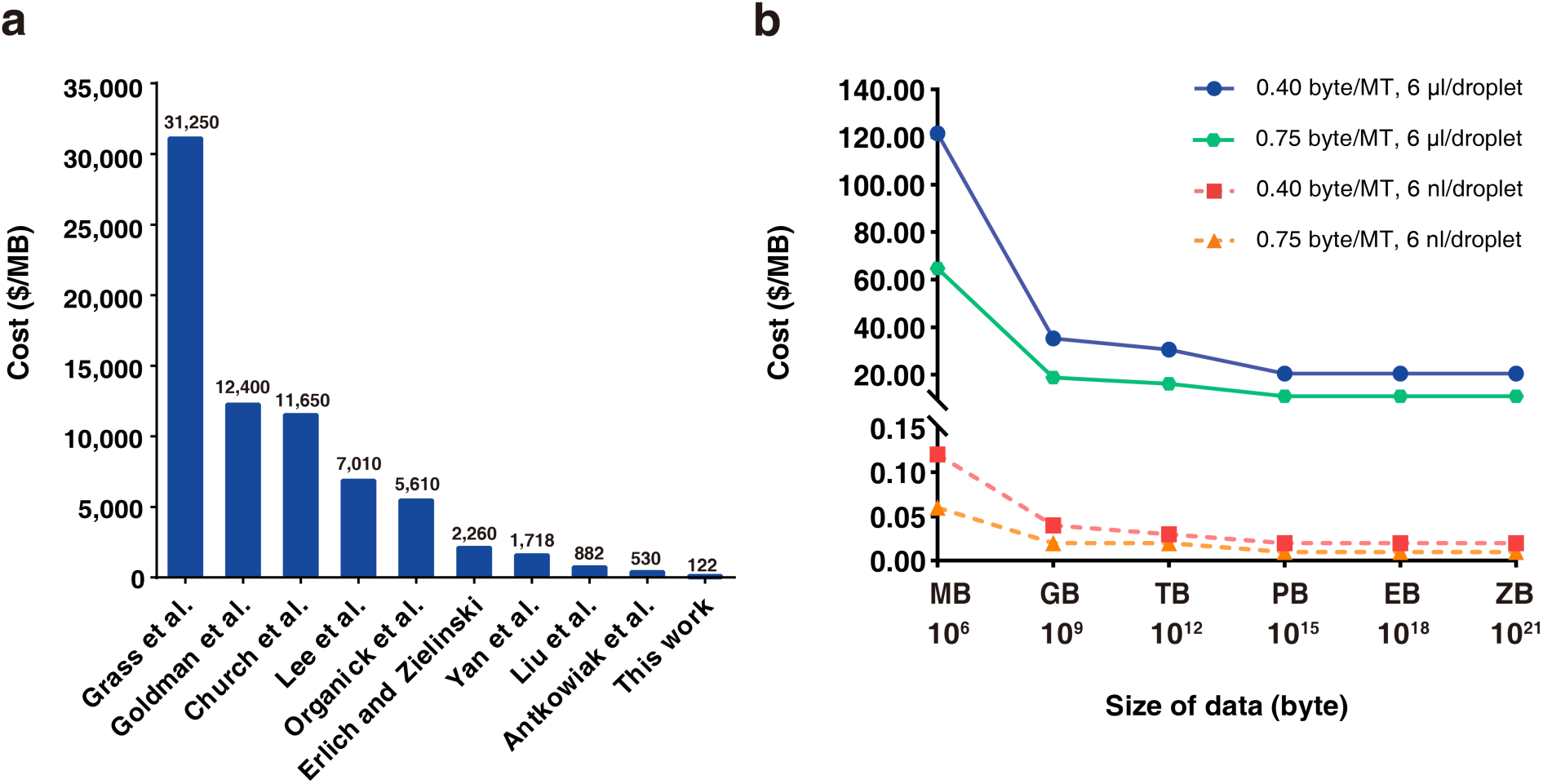
Cost-efficiency analysis of DNA-MT storage technology. a, Cost comparison between DNA-MT storage and other prevalent DNA storage technologies. Our method’s cost is estimated at 122 $/MB, in comparison to higher costs reported by other studies, such as 530 $/MB by *Antkowiak* et al., and 2,260 $/MB by *Erlich and Zielinski*. **b,** Potential for cost reduction with improvements in data size, reaction volume, and encoding density. Using the current system with 10 DNA-MTs per block and a 6 μl droplet volume, the cost of storage is 122 $ per megabyte (MB). Scaling up to store 1 gigabyte (GB) of data reduces the cost to 35.28 $/MB, and increasing to 1 terabyte (TB) further lowers it to 30.56 $/MB. By adopting a system with 20 DNA- MTs per block and maintaining a 6 μl droplet volume, the cost per MB can be reduced to 64.75 $/MB. Reducing the droplet volume to 6 nanoliters (nl) with 10 DNA-MTs per block decreases the cost per MB to 0.12 $/MB. Similarly, using 20 DNA-MTs per block with a 6 nl droplet volume further reduces the cost per MB to 0.06 $/MB. Further details on these cost calculations can be found in **Table S6**.

## 3 Discussion

Our study introduces an innovative concept in DNA storage, drawing inspiration from the ancient movable type printing to develop “DNA-MT storage”. Unlike traditional DNA storage methods that resemble block printing-simple yet costly-our system significantly reduces writing costs and time. Specifically, we have developed a comprehensive system (details in **Figure S7**) that includes DNA-MT libraries (payload, address, and checksum), a coding-decoding system (comprising DNA-MT codebooks and DNA-MT block assemblies), and an automatic DNA-MT inkjet printer, BISHENG-1. This system enables efficient storage and 100% accurate recovery of 43.7 KB diverse data, using only 350 DNA-MTs. Our approach offers a practical and scalable solution to the challenges of current DNA storage technologies.

The transformation our research brings to the DNA storage field mirrors the historical transition from block printing to movable type printing, marking a significant advance in technology and future potential.

Technologically, our innovation lies in shift from traditional DNA synthesis to enzymatic assembly of pre-made DNA-MTs, mainly offering cost and efficiency advantages.

### Cost advantage

The cost efficiency of our DNA-MT storage system is its most significant benefit over traditional DNA storage technologies. Traditional DNA storage follows a base-by-base synthesis strategy similar to block printing, necessitating the sequential encoding and synthesis of all DNA fragments from scratch. This one-time use approach precludes reusing the synthesized DNA fragments, leading to high synthesis expenses. Conversely, our system permits the repeated access and reuse of pre-made DNA-MTs, significantly reducing DNA synthesis costs. The BISHENG-1 printer showcases this cost-effective capability, where DNA-MTs, synthesized once (2 *OD*), can be used up to 10,000 times. This innovation lowers DNA storage costs to as low as 122 $/MB, given current throughput, printing speeds, and encoding density, making it the most cost-effective DNA storage solution reported thus far. [16, 20–21]

### Time advantage

Another major advantage of DNA-MT storage is its significant time efficiency. Unlike traditional methods that rely on time-consuming chemical synthesis from scratch, our approach uses biological enzyme linking, a much quicker process that enables high-throughput data writing by parallel construction of multiple DNA-MT blocks within ∼10 minutes. For example, the BISHENG-1 printer completed the writing of 43.7 KB data in ∼5 hours (4 bytes/s). In contrast, traditional DNA storage methods using chemical synthesis, such as the Dr. Oligo 192 high-throughput synthesizer (∼6 minutes per base addition, 192 channels), are estimated to take about 100 hours to synthesize the same amount of data from scratch, given the DNA storage’s capacity limit of 1.83 bits/nt. [8] This not only highlights significant time and labor savings but also enhances the practicality of DNA-MT storage.

### Targeted retrieval advantage

DNA-MT storage excels in targeted data retrieval. Each file, during the encoding and storage process, can be marked with a unique AMT for future specific retrieval. This enables the specific amplification and sequencing of all DNA-MT blocks associated with the file (employing the AMT sequence as a forward primer and a universal reverse primer from the 3’ end of the *pUC19* plasmid). This method ensures specific access to and reading of designated files, underscoring the system’s data management efficiency.

### Copy advantage

Our DNA-MT storage system enables rapid, accurate, and cost-effective data duplication through the exponential amplification of stored DNA-MT files on plasmids during bacterial culturing.

### Limitations, potential and Prospects

Since its inception in 2017, our DNA-MT storage technology has undergone several iterations. Although its current cost and efficiency are not yet suitable for commercial application, it holds considerable potential for further enhancements in both cost and time efficiency, as detailed below:

### Cost potential

There is significant untapped potential in cost reduction, including:

**i. Volume Optimization Potential**: A key improvement can be made by reducing the droplet volume used in printing each DNA-MT. Currently, the BISHENG-1 operates at a microliter scale, but transitioning to a smaller, nanoliter scale could significantly enhance cost efficiency. As demonstrated in **Figure 6b**, reducing the droplet volume to 6 nl can decrease the cost of DNA-MT storage to 0.12 $/MB. We have employed the ECHO 650 instrument (Labcyte) to decrease droplet volume to 50 nl and successfully achieved the ligation of ten DNA-MTs (**Figure S8a**), underscoring the potential for further reductions in droplet volume.
**ii. Scalability Potential:** In the current system, each DNA-MT block consists of 10 DNA-MTs. By increasing the number of DNA-MTs to 20 (4 AMTs, 15 PMTs, and 1 CMT, RS (20,10)), the storage capacity could rise to 15 bytes per block, thereby improving the encoding density from 0.40 byte per DNA-MT to 0.75 byte per DNA-MT. We have successfully achieved the ligation of 20 DNA-MTs (**Figure S5b)** and validated their assembly efficiency and accuracy in preliminary tests; however, larger-scale testing will be necessary to fully confirm these results. Assuming a 100% ligation efficiency at 0.75 byte per DNA-MT, the overall cost could be reduced by approximately 50% **(Figure 6b**). However, it is noting that expanding the library size and increasing enzyme ligation levels may introduce additional costs and could impact assembly efficiency and accuracy. Therefore, achieving effective scalability to reduce costs requires a balance of four factors: cost, the number of DNA-MTs per block, encoding density, and assembly efficiency/accuracy. On the other hand, if we reduce the droplet volume to 6 nl while simultaneously. connecting 20 DNA-MTs, the cost could be further lowered to $0.06 per MB (**Figure 6b**). Additionally, advances in encoding algorithms could further enhance both encoding density and cost-effectiveness, offering a promising direction for the future development of DNA-MT storage.
**iii. DNA-MT biosynthesis factory:** The primary cost in our current DNA-MT storage originates from DNA-MTs’ chemical pre-synthesis. Future plans involve establishing a biosynthesis factory for DNA-MTs, requiring only a single initial chemical synthesis before being inserted into plasmids for mass, cost-effective replication in bacteria. This approach would considerably lower the synthesis and production costs of pre-fabricated DNA-MTs. Furthermore, scaling and automating the biosynthesis factory could further reduce expenses, demonstrating our technology’s substantial commercial potential.

Additionally, the current cost of pre-made DNA-MT is based on commercial DNA synthesis quotations. By conducting the synthesis in-house, we could achieve substantial cost reductions. Furthermore, scaling up will spread out the expenses associated with pre-fabricated DNA-MTs and enzymes, thereby significantly lowering overall costs and amplifying savings. This strategy will render our method more economically viable.

### Efficiency potential

Our current first-gen DNA-MT injector printer, BISHENG-1, with 556 parallel channels and a 4 bytes/s speed, has significant untapped potential. We are developing the next-gen desktop DNA-MT printer, BISHENG-2, designed to substantially increase throughput and speed, aiming to enhance DNA storage efficiency and move towards industrialization and commercialization.

In summary, storage cost and efficiency are pivotal for DNA storage’s practical application. DNA-MT storage holds immense untapped potential in both aspects, awaiting further exploration and utilization. Unlocking this potential could create a more robust DNA storage ecosystem, better serving the massive data storage needs of the big data era.

## 4. Methods

### Linker candidates for DNA-MTs

To ensure high specificity and prevent unintended ligation of DNA-MTs, we selected linkers from a pool of 256 randomly generated 4-base sequences (4^4^). The selection was based on the following criteria: (1) Base diversity: Each linker must contain at least two different types of bases; (2) Homopolymer Limit: No more than three consecutive identical bases are allowed; (3) GC Content Range: 20-80%. Using these criteria, we initially screened 200 viable 4-base linker candidates.

### Linker scoring system

We randomly selected 9 linkers from linker candidate pool, supplemented by two fixed enzymatic restriction sites (*Hin*dIII: agct; *Eco*RI: aatt), yielding 55 potential linker pairs (*C*^2^_11_). A scoring system was developed to evaluate the specificity of these pairs: (1) If a linker’s sequence (forward, reverse, complementary, or reverse complementary) matches another linker’s sequence, that set of 11 linkers is discarded; (2) If a linker shares more than or equal to three consecutive or cumulative paired bases with another linker in either the forward sequence/reverse complementary sequence, and the two linkers are situated at opposite ends of the same movable type, then that combination of 11 linkers is discarded; (3) If a linker shares less than or equal to three consecutive or cumulative paired bases with another linker in either the forward sequence or reverse complementary sequence, and the two linkers are not located at opposite ends of the same movable type, then 1 point is added to this pair.

### Design of information carriers for DNA-MTs

To design effective DNA-MTs, we selected for a 20-nt length for the information carriers, balancing sequencing and synthesis costs with the need for sequence diversity. Shorter sequences lack sufficient diversity for encoding adequate information, whereas longer sequences increase both synthesis and sequencing costs. These carriers were randomly generated and coupled with the previously optimized 11 specifically optimized linkers at both 5’ and 3’ ends, generating a pool of 24/25-nt DNA-MTs (those at the ends being 25-nt in length). Several constraints were imposed on these sequences to ensure functionality and specificity: (1) Nucleotide Diversity: Each carrier must include at least three types of nucleotide bases; (2) Homopolymer Runs: Homopolymers of A/T must be less than 5, and G/C less than 4; (3) GC content: 40- 60%; (4) Restriction Site Avoidance: Exclusion of the *Eco*RI and *Hin*dIII restriction site sequences; (5) Secondary Structures: No more than 4-bp hairpin structures; (6) Self-Dimers: Consecutive self-dimers < 3, and interval self-dimers < 8; (7) Editing Distance: A minimum editing distance of six was required between any two DNA-MTs to avoid cross-hybridization.

### Construction of DNA-MT library

The single-stranded DNAs corresponding to the designed DNA-MTs were custom-synthesized and subjected to 5’-phosphorylation modifications (Shanghai Biotech Co., Ltd). These synthesized single-stranded DNA-MT oligonucleotides were then equimolarly annealed (50 μM) to form double-stranded DNA-MT solutions. Subsequently, these pre-made DNA-MTs were allocated into three sub-libraries according to the described proportions (AMT: 1,024; PMT: 1,280; and CMT: 256), ready for the assembly of DNA-MT blocks.

### Encoding target files with DNA-MTs

This section outlines the procedures for converting digital files into DNA-MT storage files, emphasizing fragmentation, data strip formation, binary conversion, error detection and correction, and final encoding into DNA-MTs in blocks. These steps ensure accurate data encoding and robust error handling to optimize the integrity of the DNA-MT stored files. **Fragmentation, strip formation, and binary conversion:** To encode various types of files (e.g., text, images, audio, and video) into DNA-MT storage files, each target file is initially segmented into 100-byte fragments. Each fragment is further subdivided into twenty 5-byte data strips. Subsequently, each strip is appended with 4 bytes of address information to facilitate mapping within the storage file, and then subjected to binary conversion. This address allocation is designed to support a total DNA-MT storage capacity of ∼2 GB. For example, as illustrated in **Figure 3** (Encoding panel) and **Figure S2**, the fourth stripe from *Shakespeare’s Sonnet 12*, “ed up…with white…do I question ma”, spanning from the 300^th^ to 399^th^ byte/character, is divided into twenty sequential 5-character data strips (from stripe 3- 0 to stripe 3-19). The first strip, 3-0, encodes the payload “ed up”, starting at the 300^th^ character, and the last strip, 3-19, encodes "on ma", beginning at the 395^th^ character. Notably, the word “white” is located in the 3-8 data strip, beginning at the 340^th^ character. Following binary conversion, the strip 3-8’s address, 340, is represented as “0, 0, 0, 98,” and its payload “white” is translated into the ASCII codes “119, 104, 105, 116, 101”. The sequence of the DNA-MT block representing “white”: agcttATCTTACAACCTCCAACCCGctccATCCCTTGCCGCTTATTCCCactaAGGGTGAAGGTGCTGATGATggtaGAAGATGTGGCGTGAGGGTGatagTCACCAATCACACCTCCTCCaagtTTTGATTCGCTGTGATGGTGctgaCGCAAGAAAGGAAAACAACGcaacAACACAAAAGGCAACACGGCaggaCGAATGGGTGGAGTGTAGGGtgtgAGTCCGTATGCCTTTTGCTGg. **Error detection and correction** (**Figure 3**: Encoding panel, and **Figure S2b**): Employing the Reed-Solomon algorithm, ten column-checksum strips (rows 20 – 29) are calculated from the twenty addressed data strips (rows 0 – 19) in each fragment. Additionally, a byte of row-checksum data, calculated using the XOR algorithm, is then appended to each of the thirty strips (twenty data strips and ten column-checksum strips). These dual-layer checksum system allows for the accurate restoration of the original data even if up to ten out of the thirty strips are compromised. **Encoding binary data into DNA-MTs:** Using the DNA-MT code table, we translate the binary data from each strip into corresponding DNA-MTs. Based on the designated DNA-MT storage codes for digital data files, these DNA-MTs are organized into 30 blocks per fragment, with each fragment encoding 100 bytes of data. Excluding ten blocks designated for checksums, each of the remaining 20 data blocks consists of four AMTs, five PMTs and one row CMT, arranged in the specified order: *A*_1_, *A*_2_, *A*_3_, *A*_4_, *P*_1_, *P*_2_, *P*_3_, *P*_4_, *P*_5_, *C*.

### Enzymatic assembly and storage of DNA-MT blocks

Following the previous DNA-MT encoding, we selected designated pre-made DNA- MTs (∼0.06 μM) from three DNA-MT libraries and distributed them into various EP tubes to construct a series of DNA-MT blocks. For each tube, along with the selected DNA-MTs, we added T4 ligase (875 U, 2011B, *TAKARA*) and 10X reaction buffer, as well as pre-digested *pUC19* modified vector (termed DNA-MT vector, 150 ng, digested with *Hin*dIII and *Eco*RI), and incubated the mixture at 25 degrees Celsius for 30 minutes, 16 degrees Celsius for 30 minutes to facilitate the sequential ligation of multiple DNA-MTs into blocks and cloned into the DNA-MT vector. Following this, the resulting DNA-MT blocks contained within the recombinant plasmids can be stored in either liquid or lyophilized form for *in vitro* storage. Furthermore, these recombinant plasmids can also be introduced into *E. coli* for *in vivo* storage. These blocks can be stored individually or combined into a single tube to form a DNA-MT stored file, representing a single digital file such as a text, image, audio, or video file.

### Modified *pUC19* plasmid

To minimize sequencing costs, the *pUC19* plasmid was modified to include Illumina sequencing adaptors and primers (**Figure S9** and **Table S7**). Specifically, we added the P5 adaptor, index 1, and Read 1 sequencing primer adjacent to the 5’ end of the *Hin*dIII restriction site. Similarly, at the 3’ end of the *Eco*RI site, the P7 adaptor, index 2, and Read 2 sequencing primer were incorporated. The modified plasmid, now termed DNA-MT plasmid, enables the direct sequencing of the DNA-MT blocks on the Illumina platform, streamlining the sequencing process.

### Design of precise spot printing of BISHENG-1

The entire printing control system of the BISHENG-1 uses a Cortex-M4 microprocessor as its core and applies mature Pulse Width Modulation (PWM) technology through a Metal-Oxide Semiconductor Field-Effect Transistor (MOS-FET) drive circuit. The MOS-FET devices are chosen for their low internal resistance and high responsiveness, making valve control more efficient. The combination of the microprocessor and the MOS-FET achieves a response time of 200 ns, and with the addition of high-precision fluidic solenoid valves, it enables accurate quantitative printing while maintaining print consistency across multiple print components within 10 μs.

### Design of intelligent monitoring of BISHENG-1

BISHENG-1 employs a high-speed industrial camera along with high-precision to monitor printing anomalies (such as missed spots or erroneous spots). AI-based image detection algorithm help achieving a detection accuracy rate of 96.69% (**Table S8**). Upon detecting any printing anomalies, the BISHENG-1 traces the error and automatically reprints. Experimental results show that the data writing efficiency of the BISHENG-1 can reach up to 4 bytes per second, with an inkjet printing accuracy rate of 99.99%.

### Preparation of printing components

Initially, the BISHENG-1 cartridges were filled with specific DNA-MTs (PMTs, AMTs, and CMTs), pre-digested DNA-MT vectors, and T4 DNA ligase, along with its reaction buffer. This preparation ensured all required components for DNA-MT printing and block assembly to ready for automated processing. To maximize the efficiency and capacity of the system, 556 cartridges were strategically allocated to optimize address space and storage density, enabling BISHENG-1 to encode 16×32^3^ blocks, equivalent to ∼2 megabytes of data with a theoretical logic density of 0.4 byte/MT. The printed file types included TXT (0.4 KB), JPEG (10.9 KB), MP3 (14.2 KB), and MP4 (18.2 KB).

### Cartridge allocation used by BISHENG-1 inkjet printing

We noticed that the simple sequential arrangement of cartridges always leads to overuse of certain cartridges and thereafter lowering their lifetime. To avoid this issue, we shuffle the cartridges by remapping the data and column-checksum strips (**Figure S10**). **AMT cartridges**: A total of 160 cartridges were distributed among the AMTs as follows: 64 cartridges for *A*_1_, including 16 cartridges for address information and 48 cartridges borrowed by payload information (**Figure S9**), and 32 cartridges each for *A*_2_, *A*_3_, and *A*_4_ . **PMT cartridges**: Each PMT from *P*_1_ to *P*_5_ was assigned 64 cartridges, totaling 320 cartridges, to ensure sufficient encoding capacity for the data blocks. **CMT cartridges**: 64 cartridges were allocated for CMTs. This allocation is based on the requirement that the row-checksum calculated from these PMTs will have 64 possible values, ensuring robust error correction capabilities. This configuration enables BISHENG-1 to cover an address space of 16 × 32^3^ blocks, resulting in a storage capacity of approximately 2 megabytes and a theoretical logic density of 0.4 byte/MT (16 × 32^3^ × 4/1024^"^).

### Loading, printing, and discharging cycle of BISHENG-1 inkjet printing

BISHENG-1 operates similarly to a standard inkjet printer but is designed to handle DNA-MTs. The process involves: **Loading**: Empty tubes are automatically loaded onto a conveyor belt, moving past various cartridges. **Printing**: As tubes pass under the designated cartridges, BISHENG-1’s control system injects the corresponding DNA- MTs, digested vectors, and ligases into the tubes. **Discharging**: Once the tubes have collected all necessary components for a DNA-MT block, they are automatically discharged from the conveyor, making room for a new cycle. This systematic approach streamlines the assembly of DNA-MT blocks. **Printing monitoring**: To address potential issues like cartridge nozzle blockages or incorrect droplet sizes, BISHENG-1 is equipped with a video camera and image analysis software. Detected errors trigger automatic reprinting of affected tubes to maintain a high success rate in DNA block assembly.

### Post-printing processing and storage

After printing, the tubes containing the DNA-MT blocks undergo enzymatic ligation at 25 degrees Celsius for 30 minutes, 16 degrees Celsius for 30 minutes. The completed blocks are then pooled together to form a DNA-MT stored file, corresponding to a digital file such as text, image, audio and video. This DNA-MT stored file can be directly stored *in vitro* or transformed into *E coli* for preservation within living organisms.

### Sequencing of the DNA-MT stored files

We initially sequenced DNA-MT stored files (stored *in vivo* or *in vitro*) to reconstruct target digital files. For *in vivo* samples, recombinant plasmids containing the stored information are first isolated and purified using a plasmid extraction kit (DP107-02, TIANGEN). Subsequently, the DNA-MT blocks within these recombinant plasmids are PCR amplified (R050Q, TAKARA) using primers Seq-F/Seq-R (details in **Table S7**). The PCR products are then purified through agarose gel electrophoresis, followed by sequencing on the Illumina NovaSeq6000 platform using PE250 paired-end sequencing technology (Novagene Inc).

### Alignment and mapping

Using the Burrows-Wheeler Alignment Tool (BWA)^44^, we aligned the DNA-MT sequences from the DNA-MT code table with paired reads, translating them into corresponding DNA-MT IDs that include four AMTs, five PMTs, and one CMT.

### Decoding and assembly

After mapping, we decoded the blocks according to the DNA-MT code table, and then converted to original information using remapping rules (**Figure S10**). This process involved translating into address and payload information from each read. If paired reads yielded conflicting results, both were discarded to ensure data integrity. Blocks were then assembled into groups of 30, with each block’s position determined by its address information. The data were reconstructed by aligning these sequences in a structured order: starting from individual strips, progressing to fragments, and finally assembling all fragments to restore the complete digital files.

Exact matches, fuzzy matches, and recombined DNA-MT sequences were identified based on voting results. Following alignment, a DNA-MT block with assigned DNA-MT IDs was generated. The exact match, fuzzy match, and recombined counts for each block were recorded along with its position and DNA-MT count in the template.

### Error correction and data recovery

Upon translating all reads, we conducted error correction by performing XOR calculations on the four AMTs and five PMTs within each block. If the XOR result matched the CMT ID, the row check passed; otherwise, the sequence was discarded as erroneous.

The output data from the row verification step are grouped based on address information into predefined sets, each consisting of 30 DNA-MT blocks. The first 20 blocks contain and store original data information, while the remaining 10 blocks are designated for redundancy column verification. If all blocks within a group are intact, column verification is then performed to identify any errors. If this verification passes, the data are confirmed for writing to the recovery file. In cases of failure, we simulate the loss of up to 10 blocks at any positions within the 30-block fragment and recalculate until the correct data is identified for writing to the recovery file. If errors persisted (for example, if there are more than 10 erroneous blocks or other unforeseen issues), that data group was recorded as erroneous in the recovery file. Finally, all payload information is written to the recovery file according to its address information (**Figure S11**).

### Storage capacity of DNA-MT storage system

The storage capacity of the system depends on its data organization and recording methods. Generally, it is calculated by multiplying the number of addressable data units the system can record and access by the capacity of each unit. This approach also applies to estimating the storage capacity of DNA-MT blocks.

To calculate the storage capacity of the current DNA-MT prototype system, we express the number of recordable and accessible addresses based on the address portion of the DNA-MT blocks:

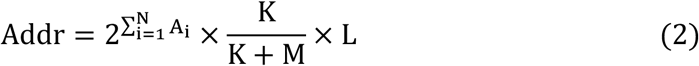

where *A*_-_ represents the bit positions of the address information for a certain level of address DNA-MTs; 𝐾 and 𝑀 are the numbers of PMTs and CMTs in the DNA-MT block, respectively; 𝐿 is the effective data payload of each DNA-MT block.

## Data availability statement

The raw data from the PE250 paired-end sequencing conducted on the Illumina NovaSeq6000 platform, have been deposited in the Genome Sequence Archive (Genomics, Proteomics & Bioinformatics, 2021) within the National Genomics Data Center (Nucleic Acids Research, 2022), hosted by the China National Center for Bioinformation / Beijing Institute of Genomics, Chinese Academy of Sciences. The datasets are publicly accessible at NGDC GSA. Specific dataset identifiers are CRA016200, corresponding to the raw data of the 4-12 DNA-MT one-step enzymatic ligation assay, and CRA016195, corresponding to the raw data of *in vitro* and *in vivo* storage of flies (TXT, JPEG, MP3, and MP4).

## Code availability statement

The code used to replicate the entire DNA movable storage process described in this study is available at ChenF Lab on GitHub. (https://github.com/ChenFLab/DNAMTStorage)

## Supporting information

Support information

## Acknowledgements

This work was supported by the National Key Research and Development Program of China (2020YFA0907000), and the National Natural Science Foundation of China (T2125013, 31971389, 32271297) for providing financial supports for this study and publication charges. The numerical calculations in this study were supported by ICT Computer-X center, and the InforSuperBahn OneAINexus platform. Chenyang Wang, Di Wei, Zheng Wei, Dongxin Yang, Jing Xing, Yunze Wang, Xiaotong Wang contributed equally to this work.

## Author contributions

Conceptualization: F.C., G.T., D.B., B.D. Methodology: C.W., D.W., Z.W., D.Y, J.X., Y.W., X.W., P.W., Writing-original draft: C.W., D.W., Z.W., D.Y, J.X., Y.W., X.W. Writing-review and editing: G.M., X.Z., H.L., C.T., P.H., J.W., R.G., G.X., C.L., Y.J.,

P.W., L.Y., Y.Z., Y.S., J.X., H.N., S.X., H.Y., D.L., F.C., G.T., D.B., B.D., Funding acquisition: F.C., D.B., G.T., Supervision: F.C., G.T., D.B., B.D.. All authors read and approved the final manuscript.

## References

[1] A. Wright, Worldwide IDC Global DataSphere Forecast, 2024–2028: AI Everywhere, But Upsurge in Data Will Take Time. International Data Corporation: 2024.

[2] a) J. Bornholt, R. Lopez, D. M. Carmean, L. Ceze, G. Seelig, K. Strauss, in Proceedings of the twenty-first international conference on architectural support for programming languages and operating systems 2016, 637–649; b) A. Extance, Nature 2016, 537 (7618), 22, 10.1038/537022a.

[3] a) L. Ceze, J. Nivala, K. Strauss, Nature Reviews Genetics 2019, 20 (8), 456, 10.1038/s41576-019-0125-3; b) M. G. T. A. Rutten, F. W. Vaandrager, J. A. A. W. Elemans, R. J. M. Nolte, Nature Reviews Chemistry 2018, 2 (11), 365, 10.1038/s41570-018-0051-5.

[4] A. Piovesan, M. C. Pelleri, F. Antonaros, P. Strippoli, M. Caracausi, L. Vitale, *BMC Res Notes* 2019, 12 (1), 106, 10.1186/s13104-019-4137-z.

[5] T. van der Valk, P. Pečnerová, D. Díez-del-Molino, A. Bergström, J. Oppenheimer, S. Hartmann, G. Xenikoudakis, J. A. Thomas, M. Dehasque, E. Sağlıcan, F. R. Fidan, I. Barnes, S. Liu, M. Somel, P. D. Heintzman, P. Nikolskiy, B. Shapiro, P. Skoglund, M. Hofreiter, A. M. Lister, A. Götherström, L. Dalén, *Nature* 2021, 591 (7849), 265, 10.1038/s41586-021-03224-9.

[6] a) E. M. LeProust, B. J. Peck, K. Spirin, H. B. McCuen, B. Moore, E. Namsaraev, M. H. Caruthers, Nucleic Acids Res 2010, 38 (8), 2522, 10.1093/nar/gkq163; b) Tucker, Tracy, Marra, Marco, Friedman, M. Jan.

[7] a) G. M. Church, Y. Gao, S. Kosuri, *Science* 2012, 337 (6102), 1628, 10.1126/science.1226355; b) N. Goldman, P. Bertone, S. Chen, C. Dessimoz, E. M. LeProust, B. Sipos, E. Birney, Nature 2013, 494 (7435), 77, 10.1038/nature11875.

[8] Y. Erlich, D. Zielinski, *Science* 2017, 355 (6328), 950, 10.1126/science.aaj2038.

[9] Z. Ping, S. Chen, G. Zhou, X. Huang, S. J. Zhu, H. Zhang, H. H. Lee, Z. Lan, J. Cui, T. Chen, W. Zhang, H. Yang, X. Xu, G. M. Church, Y. Shen, *Nat Comput Sci* 2022, 2 (4), 234, 10.1038/s43588-022-00231-2.

[10] L. Anavy, I. Vaknin, O. Atar, R. Amit, Z. Yakhini, *Nat Biotechnol* 2019, 37 (10), 1229, 10.1038/s41587-019-0240-x.

[11] W. Chen, M. Han, J. Zhou, Q. Ge, P. Wang, X. Zhang, S. Zhu, L. Song, Y. Yuan, *National Science Review* 2021, *8* (5), 10.1093/nsr/nwab028.

[12] M. Welzel, P. M. Schwarz, H. F. Löchel, T. Kabdullayeva, S. Clemens, A. Becker, B. Freisleben, D. Heider, *Nature Communications* 2023, 14 (1), 628, 10.1038/s41467-023-36297-3.

[13] C. Dou, Y. Yang, F. Zhu, B. Li, Y. Duan, *Brief Bioinform* 2024, *25* (5), 10.1093/bib/bbae363.

[14] X. Huang, J. Cui, W. Qiang, J. Ye, Y. Wang, X. Xie, Y. Li, J. Dai, *Imeta* 2024, *3* (2), e168, 10.1002/imt2.168.

[15] D. D. S. Alliance, Preserving our Digital Legacy: An Introduction to DNA Data Storage. 2021, https://dnastoragealliance.org/publications

[16] P. L. Antkowiak, J. Lietard, M. Z. Darestani, M. M. Somoza, W. J. Stark, R. Heckel, R. N. Grass, *Nat Commun* 2020, 11 (1), 5345, 10.1038/s41467-020-19148-3.

[17] a) M. Eisenstein, Nat Biotechnol 2020, 38 (10), 1113, 10.1038/s41587-020-0695-9; b) S. Palluk, D. H. Arlow, T. de Rond, S. Barthel, J. S. Kang, R. Bector, H. M. Baghdassarian, A. N. Truong, P. W. Kim, A. K. Singh, N. J. Hillson, J. D. Keasling, Nature Biotechnology 2018, 36 (7), 645, 10.1038/nbt.4173; c) H. Lee, D. J. Wiegand, K. Griswold, S. Punthambaker, H. Chun, R. E. Kohman, G. M. Church, Nat Commun 2020, 11 (1), 5246, 10.1038/s41467-020-18681-5; d) H. H. Lee, R. Kalhor, N. Goela, J. Bolot, G. M. Church, Nat Commun 2019, 10 (1), 2383, 10.1038/s41467-019-10258-1; e) J. M. Lee, J. Kwon, S. J. Lee, H. Jang, D. Kim, J. Song, K. T. Kim, Sci Adv 2022, 8 (10), eabl8614, 10.1126/sciadv.abl8614; f) B. H. Nguyen, C. N. Takahashi, G. Gupta, J. A. Smith, R. Rouse, P. Berndt, S. Yekhanin, D. P. Ward, S. D. Ang, P. Garvan, H. Y. Parker, R. Carlson, D. Carmean, L. Ceze, K. Strauss, Sci Adv 2021, 7 (48), eabi6714, 10.1126/sciadv.abi6714.

[18] a) Chen, F., Bu, D., Ma, G., Wang, C. & Xing, J., China patent: CN202010688281.X, 2020; b) Chen, F., Bu, D., Ma, G., Wang, C. & Xing, J., United States patent: US20230274793A1, 2021.

[19] Z.Y. Gong, L.F. Song, G.S. Pei, Y.F. Dong, B.-Z. Li, Y.J. Yuan, *Engineering* 2023, 29, 130, 10.1016/j.eng.2022.05.023.

[20] C. Xu, B. Ma, X. Dong, L. Lei, Q. Hao, C. Zhao, H. Liu, ACS Applied Materials & Interfaces 2023, 15 (20), 24097, 10.1021/acsami.3c01860.

[21] C. Zhang, R. Wu, F. Sun, Y. Lin, Y. Liang, J. Teng, N. Liu, Q. Ouyang, L. Qian, H. Yan, *Nature* 2024, 634 (8035), 824, 10.1038/s41586-024-08040-5.

[22] a) J. Xing et al., China patent: CN202010381206.9, 2020; b) B. Duan et al., China patent: CN202111045564.3, 2021; c) B. Duan et al., China patent: CN202110976951.2, 2021; d) B. Duan. et al., China patent: CN202111044054.4, 2021; e) B. Duan et al., China patent: CN202110975074.7, 2021; f) B. Duan et al., China patent: CN202111045576.6, 2021; g) B. Duan et al., China patent: CN202111045562.4, 2021.; h) B. Duan et al., China patent: CN202111044027.7, 2021; i) B. Duan et al., China patent: CN202111045588.9, 2021; j) B. Duan et al., China patent: CN 202111465019.X, 2021.

[23] J. S. Plank, Software: Practice and Experience 1997, 27 (9), 995.

[24] D. A. Patterson, G. Gibson, R. H. Katz, presented at Proceedings of the 1988 ACM SIGMOD international conference on Management of data, Chicago, Illinois, USA, 1988.

[25] S. Shin, J. Park, Mol. BioSyst. 2016, 12, 10.1039/C5MB00750J.

[26] G. Navarro, ACM Comput. Surv. 2001, 33 (1), 31, 10.1145/375360.375365.

[27] N. C. o. t. I. U. o. B. (NC-IUB), European Journal of Biochemistry 1979, 97 (2), 319, 10.1111/j.1432-1033.1979.tb13116.x.

