## Supplementary material for "Cost-effective DNA Storage System with DNA Movable Type": Support information

| **File format** | **#Blocks number** | **Size**  **(bytes)** | **Note** |
| --- | --- | --- | --- |
| TXT | 180 | 434 | Chinese Tang Poetry  “*SHUI DIAO GE TOU*” |
| JPEG | 4200 | 11,128 | Picture of “*PANDA*” |
| MP3 | 5460 | 14,490 | The song "*QI CHENG*" |
| MP4 | 6990 | 18,605 | The video “*Horse*” |

Footnotes: “*Horse*” adapted from Eadweard Muybridge, Human and Animal Locomotion, plate 626, thoroughbred bay mare ‘Annie G.’ galloping, Wikimedia Commons.

**Table S1. Details of four printed DNA movable type stored files using BISHENG-1.** This table provides information on four files stored using DNA-MTs, including their format, size, number of DNA-MT blocks required, and a brief description of the content.

| **ID.** | **Sequence (5’-3’)** |
| --- | --- |
| 1 | **agctt**ATACTCCTTTCTTCGGTCGG**ctcc**ATCCCTTGCCGCTTATTCCC**acta**AGGGTGAAGGTGCTGATGAT**ggta**GCAATGAGGAACAACAGACG**atag**CAAACCGCCCAAAACCAACG**aagt**TTTCTATGTGCTCTCCGTCT**ctga**GAACACCCACCCAAAGAGGA**caac**AGCAAACTACATCGGACAAC**agga**TTTAGGTTGGTGGGAGAGGT**tgtg**CTGGGTAGTTTGGGTGCCGT**gaattc** |
| 2 | **agctt**ATCTTACAACCTCCAACCCGctccTCCCGCATCCCTCTGTTCCT**acta**TGAACAAATGGAGGAGGCGA**ggta**GGGTTGGAGAAAGGGATAGG**atag**ACAGGGAACAAGGAGTGCGA**aagt**AGTATTGTGAGTGGGCGTGG**ctga**GCATACTTCCCTTCCACCTA**caac**ACAATACGCAGAAAGCCAGA**agga**TTTTATTCGGCAGTGGGCTT**tgtg**AGGGAGAGTGGTGGCATCGT**gattc** |
| 3 | **agctt**CGTCTCCCAATAATCCAGTC**ctcc**CTTCCTCGCTCTACCTGTCC**acta**TTTGGTTCTTCTTGGCGGTG**ggta**GAGAGCAGGAAATAATAGGG**atag**GCGTAAAAGGAGAGGAGTCA**aagt**AACACCACCGCCCACCAAAT**ctga**ACTACCAAGCGACCACCCAC**caac**AGGAGTGAAACGACGAAACG**agga**GGGAGTGTTTATGGGCGAGT**tgtg**AGATTGGAGGGCGGTATTTG**gattc** |
| 4 | **agctt**TCTTCCAGCGTAGTGTCGGT**ctcc**GACCCTTTTCACCTCCAGCC**acta**AGCAAGGCGAGAAACAAAAG**ggta**GGCGTTGTAGTGATGTTGGG**atag**GGATGGGCGGTGAGAAAGAA**aagt**GTGAGCAAAAGGAAGCGAGT**ctga**GCCTACCAACGACAACGAGA**caac**ACCGCAAGGATAAGGACCGA**agga**GAGGAGGTGTTGAGTTTGAT**tgtg**CCCTCCGTTGCTATCTTACT**gattc** |
| 5 | **agctt**CCGTTGTATTCTCTCGTCCGctccTTACTCCCTACAAATCGCAC**acta**TTGTTTTCCTCCTTGTCCTG**ggta**AGAAGCAGCCCATAGAGGAA**atag**AAATACGACCCAACACAGAC**aagt**GTTCCGATGTTGAGGTGGTG**ctga**CACCGCCCTACAACTGAAAA**caac**CACGGACAAGAAGCATCAAA**agga**AGTAGGGTTGTGCGGGTGAA**tgtg**CTAAGAGGTTGGAGGGTGCT**gattc** |
| 6 | **agctt**GATACCCTCCACGACCAATActccGTTCTTACCATCTTTCCCTG**acta**AGACAAGAACAGAGAAGCCA**ggta**GGCAAGATGAAAGGCTCGGA**atag**CGGAACCAGACGAAGGAGAA**aagt**GGAGGATTGGTTGGTTGCGT**ctga**CCACGGGAAACTAAACCACA**caac**AGCAGAGGAGGTCAAAGTCA**agga**AGTGAATGTTTGGGAGGTGT**tgtg**TGTTTGGTTAGGGAGCGATG**gattc** |
| 7 | **agctt**ACTGTCTAATGGTCGTGGTTctccATCTCGCCGCTCAACACTTT**acta**TGGGAGGGACGGGATTTATG**ggta**GAGTTTCGTGGGTTTGGGAC**atag**CGGGTCGTGAGGAGAATAAT**aagt**CTCCTGGTTGTATGGCTGTG**ctga**GAACCAACCTTACCGACAAC**caac**CACTAAAGGAAAGCAGCCCA**agga**AGTGGTGAGAAGAGCAGGGA**tgtg**TGAGTCCATCCTGCTGTTGA**gattc** |
| 8 | **agctt**TTGCCTCGTCTGAACTTGTG**ctcc**ACCCACTCAACTATGCTATC**acta**ATGAGAAAGGAGGGTGGGCG**ggta**GTGGGAAGACAAATGGTGGG**atag**CGAGCAGAGAAGACGACAAC**aagt**GTAGAGGGTTGGTATTGTGG**ctga**ATCCAAGACAAAACCGCTGA**caac**GAGAGTGCGACCATAACAAA**agga**GGCGGGTGTGTAGTATCAGT**tgtg**TTTGGTGGTCCCGATTGTTT**gattc** |

Footnotes: Lowercase bases represent linkers (4 nt), and uppercase bases represent information carriers (20 bp).

**Table S2.** **DNA-MT blocks with 52-score linker group.**

| **ID.** | **Sequence (5’-3’)** |
| --- | --- |
| 1 | **aagctt**ATACTCCTTTCTTCGGTCGG**ctcc**CGTTTGCTCCGTTTCCTGTT**atgc**TTGCTGCTTCGCTTTCTCTG**ggta**GCAATGAGGAACAACAGACG**atag**ATGGAAGAAATGATAGGCGG**agac**GGCGAAGAGAGTAGATAGTG**ctga**GAACACCCACCCAAAGAGGA**caac**GGCAAGCACCAGAACCCAAC**acga**GTAGGAGCAAGGTAAAGCGG**tgtg**CTGGGTAGTTTGGGTGCCGT**gaattc** |
| 2 | **aagctt**ATCTTACAACCTCCAACCCG**ctcc**CGTTCCAATCCACCAAATCT**atgc**TCGGTTGTGTTTGTAGGCTC**ggta**GGGTTGGAGAAAGGGATAGG**atag**CATAGGAAAGGCGGAAAGGG**agac**CCTCCCTAAAACAACCCGCC**ctga**GCATACTTCCCTTCCACCTA**caac**ACACGGCAACCTCTCACTCC**acga**CGCAAATAGAAGGAAAGGAC**tgtg**AGGGAGAGTGGTGGCATCGT**gaattc** |
| 3 | **aagctt**CGTCTCCCAATAATCCAGTC**ctcc**TTGTTACCTCTCCTTGCCGA**atgc**TCTGTGGTTCTTATGGTGGT**ggta**GAGAGCAGGAAATAATAGGG**atag**AACGAGAGGGAGGGCATAAG**agac**AATCCCATACGAACCAAGAC**ctga**ACTACCAAGCGACCACCCAC**caac**GCACCACAGAATAGCCCACC**acga**ATAGGGAGAAGAAGGTGGAA**tgtg**AGATTGGAGGGCGGTATTTG**gaattc** |
| 4 | **aagctt**TCTTCCAGCGTAGTGTCGGT**ctcc**AAGAACTCAACACCCACACT**atgc**TCTTGGGATGTTGGGTTTCG**ggta**GGCGTTGTAGTGATGTTGGG**atag**CATAAAGGAAAACACGGCAC**agac**CCGCCGCAAGAAGACCAACA**ctga**GCCTACCAACGACAACGAGA**caac**CAGGAAAAGGGCAAGAACAT**acga**TTGGTGAGGTGGAGGGTAGA**tgtg**CCCTCCGTTGCTATCTTACT**gaattc** |
| 5 | **aagctt**CCGTTGTATTCTCTCGTCCG**ctcc**GTTTCCCTGTTTTGTCCCTA**atgc**GGTGGGCGATTTAGTTGGGA**ggta**AGAAGCAGCCCATAGAGGAA**atag**CAAGAATAAGAACGACCAGC**agac**CCACGCCTACGCAAAACAAA**ctga**CACCGCCCTACAACTGAAAA**caac**AGAACAGACAGCGACAGAAT**acga**GCCAGAAGAAAAGAACGCAG**tgtg**CTAAGAGGTTGGAGGGTGCT**gaattc** |
| 6 | **aagctt**GATACCCTCCACGACCAATA**ctcc**GCTATCCCACCTATCACCTT**atgc**CGAGGTGGGTGGGTGCTAAA**ggta**GGCAAGATGAAAGGCTCGGA**atag**ACACGCCAAAGAACAGAGGT**agac**GAAGGAGACGAACTGCCGAA**ctga**CCACGGGAAACTAAACCACA**caac**ACAAAAGAGCGAGAGTCCGA**acga**GTCCCTTCTTTTCCCTTGCC**tgtg**TGTTTGGTTAGGGAGCGATG**gaattc** |
| 7 | **aagctt**ACTGTCTAATGGTCGTGGTT**ctcc**TCCTCAACCCACCCAATCAA**atgc**TGCGGGCTGTGGTTATTTGT**ggta**GAGTTTCGTGGGTTTGGGAC**atag**AAACGAGCAAACCCAACCAG**agac**GACTACACCCAAATCCCTAA**ctga**GAACCAACCTTACCGACAAC**caac**ACCCAGTAGCAAGAAAGACA**acga**TGTGGTGGGAGGTTAGGTTA**tgtg**TGAGTCCATCCTGCTGTTGA**gaattc** |
| 8 | **aagctt**TTGCCTCGTCTGAACTTGTG**ctcc**TTGTCCACTCCACCCTCCTA**atgc**TCCCGCCTCGTTCTGCTTTT**ggta**GTGGGAAGACAAATGGTGGG**atag**CCAAAGAATAGCACAAGAGG**agac**GCAAACTCCCAAACAAAGAC**ctga**ATCCAAGACAAAACCGCTGA**caac**CACCCAGCGGACAAACGATA**acga**TTGTGGGTAGGTGGTGGCTT**tgtg**TTTGGTGGTCCCGATTGTTT**gaattc** |

Footnotes: Lowercase bases represent linkers (4 nt), and uppercase bases represent information carriers (20 bp).

**Table S3.** **DNA-MT blocks with 55-score linker group.**

| **ID.** | **Sequence (5’-3’)** |
| --- | --- |
| 1 | **aagctt**TTCCACCTCACTCATTCTCC**ctcc**AAACCTCCTCCTCCCTACTT**acta**TTTGGGTGGTGTTTGTGGAG**ggta**AGAAGAGGAGACGAGCGATA**gaattc** |
| 2 | **aagctt**TTCCACCTCACTCATTCTCC**ctcc**AAACCTCCTCCTCCCTACTT**acta**AGGGAAGAGAAGAAGGTGGA**ggta**GTGGATGGTTAGGTTGTGTC**gaattc** |
| 3 | **aagctt**TTCCACCTCACTCATTCTCC**ctcc**AAACCTCCTCCTCCCTACTT**acta**AAGGAAGGAGGAGAGGACAA**ggta**GAGTGAGGTAGAAGAGGTGA**gaattc** |
| 4 | **aagctt**TTCCACCTCACTCATTCTCC**ctcc**AAACCTCCTCCTCCCTACTT**acta**AGGAAAGGAGGGACAGAACA**ggta**GAGGGAAGTAGGATGATGGT**gaattc** |
| 5 | **aagctt**TTCCACCTCACTCATTCTCC**ctcc**AAACCTCCTCCTCCCTACTT**acta**TTGTTGGGTGTGAGGTGTCT**ggta**GGAGGGTTATGTGGTTTGGT**gaattc** |
| 6 | **aagctt**TTCCACCTCACTCATTCTCC**ctcc**AAACCTCCTCCTCCCTACTT**acta**TGGGTGGTTTGTGGTTGAAG**ggta**TTTGGTTGGGTGTGCCGTTA**gaattc** |
| 7 | **aagctt**TTCCACCTCACTCATTCTCC**ctcc**AAACCTCCTCCTCCCTACTT**acta**TGTTTGGTGGTTGTCGCTCT**ggta**AGGAGGAGATGGATGAGGAA**gaattc** |
| 8 | **aagctt**TTCCACCTCACTCATTCTCC**ctcc**AAACCTCCTCCTCCCTACTT**acta**TGGTTTGGGTGTGTTGGTGT**ggta**AGGGTGAAGATGTGAGAGGA**gaattc** |
| 9 | **aagctt**TTCCACCTCACTCATTCTCC**ctcc**AAACCTCCTCCTCCCTACTT**acta**TGTGTGTTGGGTTTGGTTCG**ggta**TTGCCTTGTCTCCCGTGTTA**gaattc** |
| 10 | **aagctt**TTCCACCTCACTCATTCTCC**ctcc**AAACCTCCTCCTCCCTACTT**acta**TGTGGGTGTTGTGGGTGTAA**ggta**GTGTATGAGCGGGAGGTTTA**gaattc** |
| 11 | **aagctt**TTCCACCTCACTCATTCTCC**ctcc**AAACCTCCTCCTCCCTACTT**acta**TGTGTGTGTGGTGTGTAAGC**ggta**TTTTGGGTGGTGGAGCGTTA**gaattc** |
| 12 | **aagctt**TTCCACCTCACTCATTCTCC**ctcc**AAACCTCCTCCTCCCTACTT**acta**AAGAAGGGAGGAGGAAAAGC**ggta**AAGTGAGAGCGGGAGAAGAT**gaattc** |

Footnotes: Lowercase bases represent linkers (4 nt), and uppercase bases represent information carriers (20 bp).

**Table S4.** **DNA-MT blocks for preliminary carrier design experiment (20 bp).**

| **ID.** | **Sequence (5’-3’)** |
| --- | --- |
| 1 | **aagctt**GTTGTTCCTCTCCCTTCTCTTTCCACTCTC**ctcc**TCCAACACCAACCAACCCAACAACCACTACactaTTGGGTTGTGGTGGTGTTGTGTGTTGTTCG**ggta**AAAGGAGAGGGAGACAGAAGGAAAAGAGGC**gaattc** |
| 2 | **aagctt**GTTGTTCCTCTCCCTTCTCTTTCCACTCTC**ctcc**TCCAACACCAACCAACCCAACAACCACTAC**acta**TTGTGGCTCTGGTTTTGGTCGTGTGTGTTC**ggta**AGCAGAAACGAACACGGAGAAAGACAGGAG**gaattc** |
| 3 | **aagctt**GTTGTTCCTCTCCCTTCTCTTTCCACTCTC**ctcc**TCCAACACCAACCAACCCAACAACCACTAC**acta**TTTGTGGGTGGTATCGGTTGTGGGCTGTTT**ggta**TGTTGTAGGGTTGGAGGTGATGAGGCGTTT**gaattc** |
| 4 | **aagctt**GTTGTTCCTCTCCCTTCTCTTTCCACTCTC**ctcc**TCCAACACCAACCAACCCAACAACCACTAC**acta**TTGGTGTGTGTATCGTGTGGGTTTTGGGTC**ggta**AAGGGTGTGTTGGTTGGGTTGGCTGTGATT**gaattc** |
| 5 | **aagctt**GTTGTTCCTCTCCCTTCTCTTTCCACTCTC**ctcc**TCCAACACCAACCAACCCAACAACCACTAC**acta**TTTGCTGGTCTGGTTTGGTCGTGTGGTCTT**ggta**GGTGAGTTTGTTGGTTGGAGTTGGGAGTGA**gaattc** |
| 6 | **aagctt**GTTGTTCCTCTCCCTTCTCTTTCCACTCTC**ctcc**TCCAACACCAACCAACCCAACAACCACTAC**acta**AAGGACGAGAGAAAGAAGGGTGAGGAGATG**ggta**AAGAGCAGCAGCAAGAAGACAAAGGGCAAC**gaattc** |
| 7 | **aagctt**GTTGTTCCTCTCCCTTCTCTTTCCACTCTC**ctcc**TCCAACACCAACCAACCCAACAACCACTAC**acta**TGTGGCTTCTGTTGGGTTTGCTGGTGTTGT**ggta**AATGGAGTGGTAGGTAGGAGGGATAGGGTT**gaattc** |
| 8 | **aagctt**GTTGTTCCTCTCCCTTCTCTTTCCACTCTC**ctcc**TCCAACACCAACCAACCCAACAACCACTAC**acta**TGTTTGGGTGTTCTTGTGGGTGTGCTTGCT**ggta**AGGATTGGGAGGATGAAGGAGTTTGGGATG**gaattc** |
| 9 | **aagctt**GTTGTTCCTCTCCCTTCTCTTTCCACTCTC**ctcc**TCCAACACCAACCAACCCAACAACCACTAC**acta**TTCTGTTGCGTGTGTGGTTGTTCGTTGGTC**ggta**GTGGGAAAGTGGTATGAGAGAGGAGAGGAA**gaattc** |
| 10 | **aagctt**GTTGTTCCTCTCCCTTCTCTTTCCACTCTC**ctcc**TCCAACACCAACCAACCCAACAACCACTAC**acta**TTGTTCTGGCGGTGTTTGTGTGTGTCGTTC**ggta**TGGGAGTTGAGCGAAAGGTTGTGGAGTGTA**gaattc** |
| 11 | **aagctt**GTTGTTCCTCTCCCTTCTCTTTCCACTCTC**ctcc**TCCAACACCAACCAACCCAACAACCACTAC**acta**TGGAGGTTTGTGTTGGTGGATGTGTGGTCT**ggta**GTGTTGAGGAAGTAGAGGAGAGGGAGGATA**gaattc** |
| 12 | **aagctt**GTTGTTCCTCTCCCTTCTCTTTCCACTCTC**ctcc**TCCAACACCAACCAACCCAACAACCACTAC**acta**TGGTGTGTTGGTTTTGTGCGATTTGTGGGC**ggta**GTTTTGCCTGCGTTTGGTGTTCGGTTTGTG**gaattc** |

Footnotes: Lowercase bases represent linkers (4 nt), and uppercase bases represent information carriers (30 bp).

**Table S5. DNA-MT blocks for preliminary carrier design experiment (30 bp).**

| **Scheme** | **Data size** | ***S***  **(US$)** | ***V***  **(ul)** | $\boldsymbol{\rho}$ **(byte/ul)** | ***L***  **(US$/byte)** | ***C* (US$/MB)** |
| --- | --- | --- | --- | --- | --- | --- |
| 6 μl/droplet 0.4byte/MT | MB | 1.21E+02 | 2.36E+07 | 0.04 | 2.19E-07 | 121.57 |
|  | GB | 3.51E+04 | 2.36E+10 | 0.04 | 2.19E-07 | 35.28 |
|  | TB | 3.03E+07 | 2.36E+13 | 0.04 | 2.19E-07 | 30.56 |
|  | PB | 2.02E+10 | 2.36E+16 | 0.04 | 2.19E-07 | 20.45 |
|  | EB | 2.02E+13 | 2.36E+19 | 0.04 | 2.19E-07 | 20.45 |
|  | ZB | 2.02E+16 | 2.36E+22 | 0.04 | 2.19E-07 | 20.45 |
| 6 μl/droplet 0.75byte/MT | MB | 6.48E+01 | 1.26E+07 | 0.08 | 3.81E-08 | 64.75 |
|  | GB | 1.87E+04 | 1.26E+10 | 0.08 | 3.81E-08 | 18.73 |
|  | TB | 1.62E+07 | 1.26E+13 | 0.08 | 3.81E-08 | 16.22 |
|  | PB | 1.08E+10 | 1.26E+16 | 0.08 | 3.81E-08 | 10.83 |
|  | EB | 1.08E+13 | 1.26E+19 | 0.08 | 3.81E-08 | 10.83 |
|  | ZB | 1.08E+16 | 1.26E+22 | 0.08 | 3.81E-08 | 10.83 |
| 6 nl/droplet 0.4byte/MT | MB | 1.29E-01 | 2.50E+04 | 44.44 | 2.19E-10 | 0.12 |
|  | GB | 3.71E+01 | 2.50E+07 | 44.44 | 2.19E-10 | 0.04 |
|  | TB | 3.21E+04 | 2.50E+10 | 44.44 | 2.19E-10 | 0.03 |
|  | PB | 2.14E+07 | 2.50E+13 | 44.44 | 2.19E-10 | 0.02 |
|  | EB | 2.14E+10 | 2.50E+16 | 44.44 | 2.19E-10 | 0.02 |
|  | ZB | 2.14E+13 | 2.50E+19 | 44.44 | 2.19E-10 | 0.02 |
| 6 nl/droplet 0.75byte/MT | MB | 7.71E-02 | 1.50E+04 | 83.33 | 3.81E-11 | 0.06 |
|  | GB | 2.23E+01 | 1.50E+07 | 83.33 | 3.81E-11 | 0.02 |
|  | TB | 1.93E+04 | 1.50E+10 | 83.33 | 3.81E-11 | 0.02 |
|  | PB | 1.29E+07 | 1.50E+13 | 83.33 | 3.81E-11 | 0.01 |
|  | EB | 1.29E+10 | 1.50E+16 | 83.33 | 3.81E-11 | 0.01 |
|  | ZB | 1.29E+13 | 1.50E+19 | 83.33 | 3.81E-11 | 0.01 |

**Table S6.** **Cost calculation for DNA movable type storage.** This table provides detailed cost calculations related to Figure 6b, showing cost reductions with increases in reaction volume, encoding density and data size improvements.

| **Name** | **Sequence (5’-3’)** | **Function** |
| --- | --- | --- |
| Test-F | CATTAGGCACCCCAGGCTT | Plasmid identification & Sanger sequencing |
| Test-R | GCACAGATGCGTAAGGAGAAAAT |  |
| Seq-F | AATGATACGGCGACCACC | Next-generation sequencing library construction |
| Seq-R | CAAGCAGAAGACGGCATACG |  |
| HF | AGCTAGCTT*AATGATACGGCGACCACCGAGATCTACAC***AGTGGTCA**TCGTCGGCAGCGTCAGATGTGTATAAGAGACAGA | Insert fragment  (P5 adaptor + index1 + read1 sequencing primer) |
| HR | AGCTTCTGTCTCTTATACACATCTGACGCTGCCGACGA**TGACCACT***GTGTAGATCTCGGTGGTCGCCGTATCATT*AAGCT |  |
| EF | AATTC*CTGTCTCTTATACACATCTCCGAGCCCACGAGAC***TATCAGCA**ATCTCGTATGCCGTCTTCTGCTTGGAATT | Insert fragment  (P7 adaptor + index2 + read2 sequencing primer) |
| ER | AATTAATTCCAAGCAGAAGACGGCATACGAGAT**TGCTGATA***GTCTCGTGGGCTCGGAGATGTGTATAAGAGACAG*G |  |

Footnotes: In the sequences, italic indicates the adaptor sequence, bold indicates the index sequence, and underlined text denotes the read sequencing primer sequence.

**Table S7.** **Primer sequences for constructing DNA movable type block storage plasmids (Figure S9).** The table lists the primer sequences used to construct DNA-MT plasmids. HF + HR primers insert the P5 adaptor, index 1, and Read 1 sequencing primer at the 5’ end of the HindIII site. EF + ER primers add the P7 adaptor, index 2, and Read 2 sequencing primer at the 3' end of the *Eco*RI site. Test-F/Test-R primers are used to amplify DNA-MT blocks in recombinant plasmids for Sanger sequencing. Seq-F/Seqt-R primers are used for PCR amplification of DNA-MT blocks in recombinant plasmids, enabling the direct construction of next-generation sequencing libraries for DNA-MT blocks.

| Resnet num^a)^ | Learning rate | Epoch^b)^ | mAP^c)^ | Infer time(ms)^d)^ |
| --- | --- | --- | --- | --- |
| 22  56 | 1e-4  1e-4 | 1000  1000 | 82.61%  90.16% | 2.44  4.35 |
| 110  152 | 1e-4  1e-4 | 1000  1000 | 96.09%  96.69% | 7.29  8.51 |

Footnotes: ^a)^Number of layers in the ResNet model; ^b)^Number of training epochs; ^c)^Mean average precision; ^d)^Inference time;

**Table S8. Training results of deep learning-based object detection algorithm for the BISHENG-1 system.**

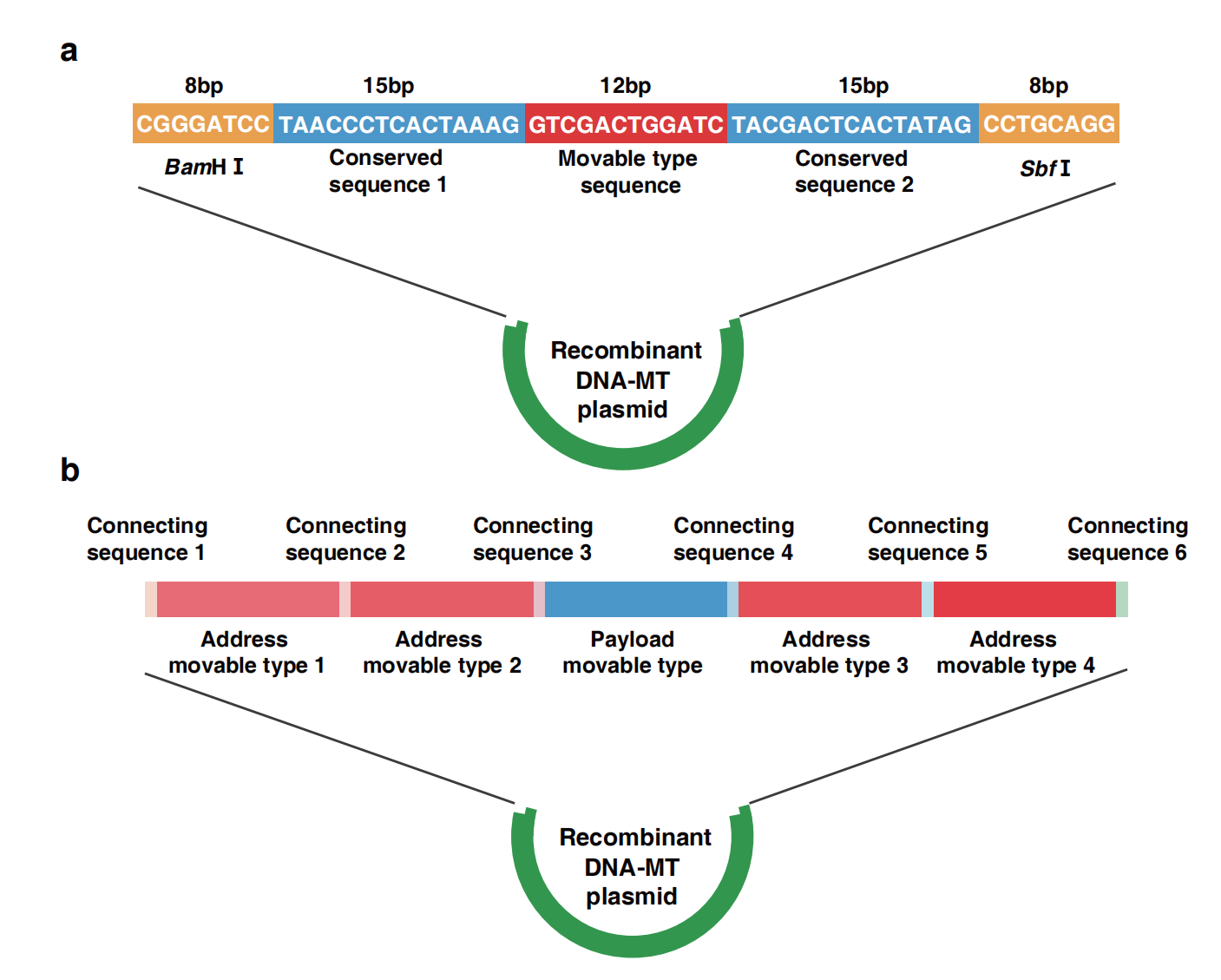

**Figure S1. First-generation DNA movable type structure. a, The first-generation DNA movable type structure**. Adapted from page 10 of the grant application to the National Natural Science Foundation of China in 2019 (Grant Title: A Study on a Novel DNA Storage Technology, Grant No.: 31971389, Duration: 2020.1-2023.12). **b, One-step (or multi-step) PCR amplification construction of DNA movable type blocks for cloning into the pUC19 plasmid.** Adapted from page 51 of the Chinese patent (202010688281.X, dated 2020.07.16) and page 2 of the PCT/US patents (PCT/CN2021/098663, dated 2021.07.07, and US20230274793A1, dated 2023.01.13).

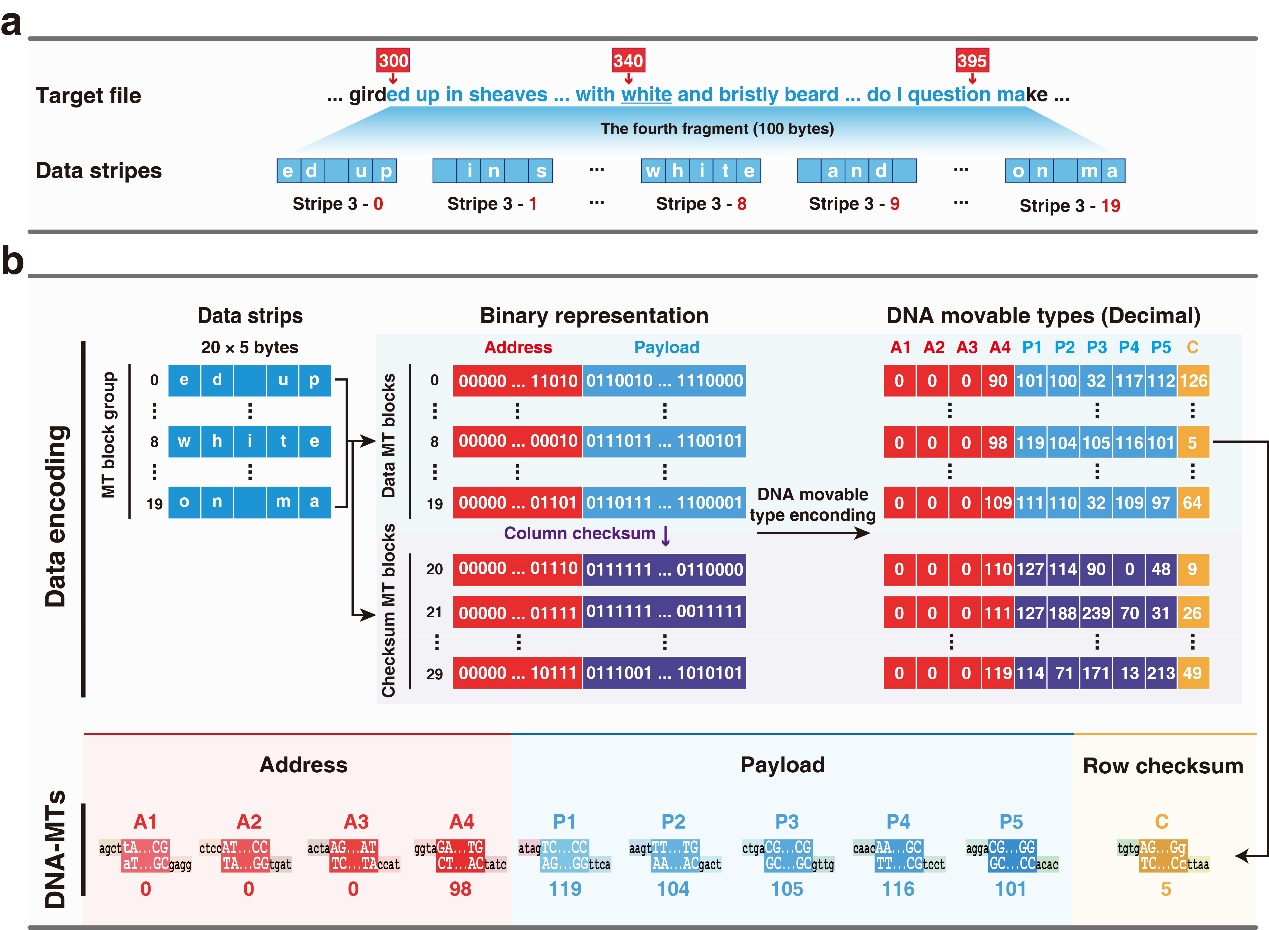

**Figure S2.** **Conversion of target files into DNA-MT blocks. a, File fragmentation and data strip formation.** The target file is divided into 100-byte/character fragments. As illustrated, the fragment “ed up...with white...do I question ma” (spanning characters 300 to 399) is subdivided into twenty sequential 5-character data strips. The first strip begins with the characters “ed up” at position 300, while the last strip encodes “on ma” beginning at position 395. The word “white” is highlighted in the 8^th^ data strip, beginning at character 340. **b, Conversion to DNA-MT blocks.** Each of the twenty data strips (rows 0-19) is appended with four-byte address information to form addressed data strips, which is further translated into binary representations. For error detection and correction, ten column-checksum strips (rows 20 – 29) and a row-checksum DNA-MT (column 9) are incorporated. Utilizing a code table, characters from the strips are mapped to their corresponding DNA-MTs, leading to the assembly of DNA-MT blocks. Specifically, the DNA-MT block encoding the word “white” in the 8^th^ strip comprises four AMTs ($A_{1}$-0, $A_{2}$-0, $A_{3}$-0, $A_{4}$-98), five PMTs ($P_{1}$-119, $P_{2}$-104, $P_{3}$-105, $P_{4}$-116, $P_{5}$-101) and one CMT ($C$-5).

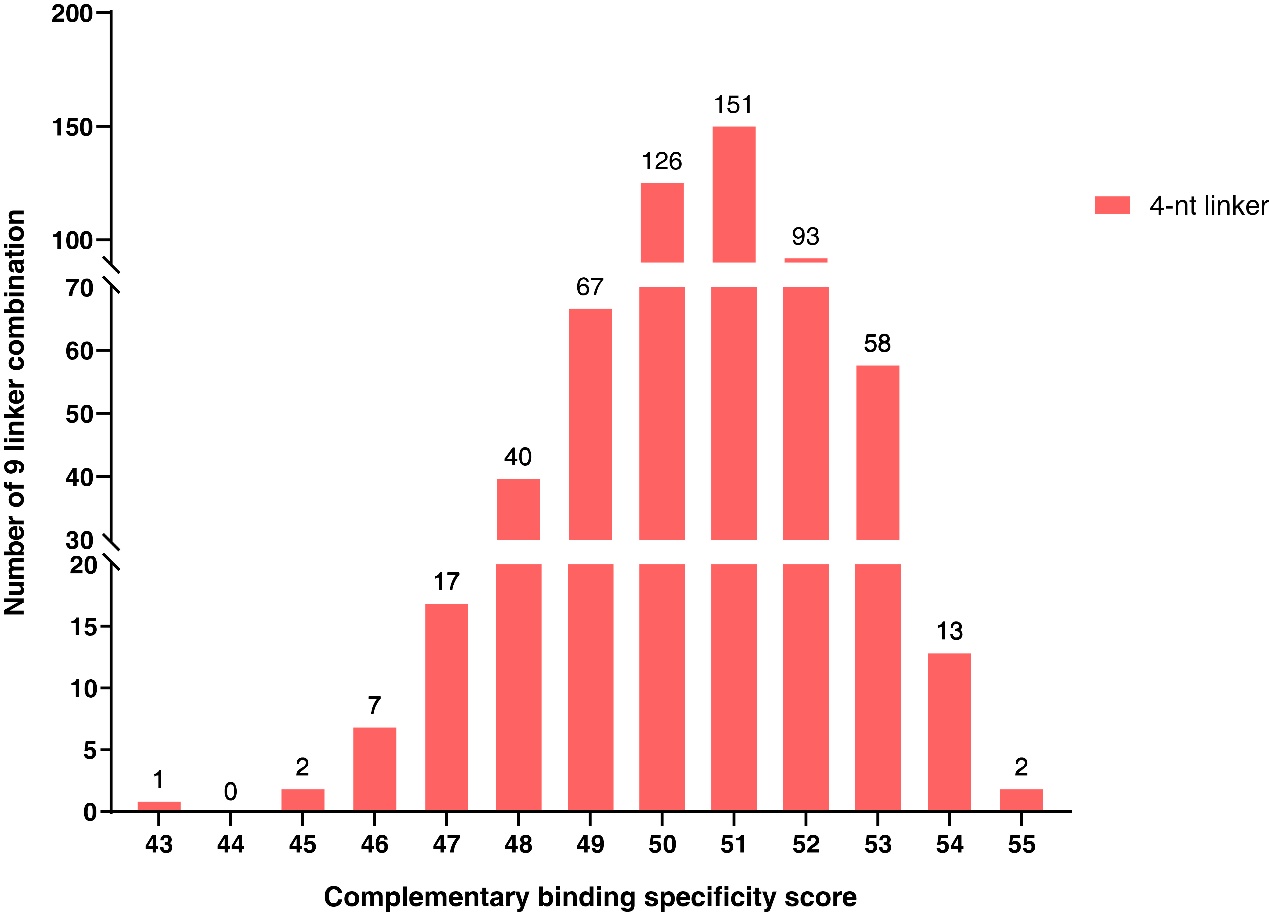

**Figure S3.** **Complementary binding specificity scores of 9 linker combinations.**

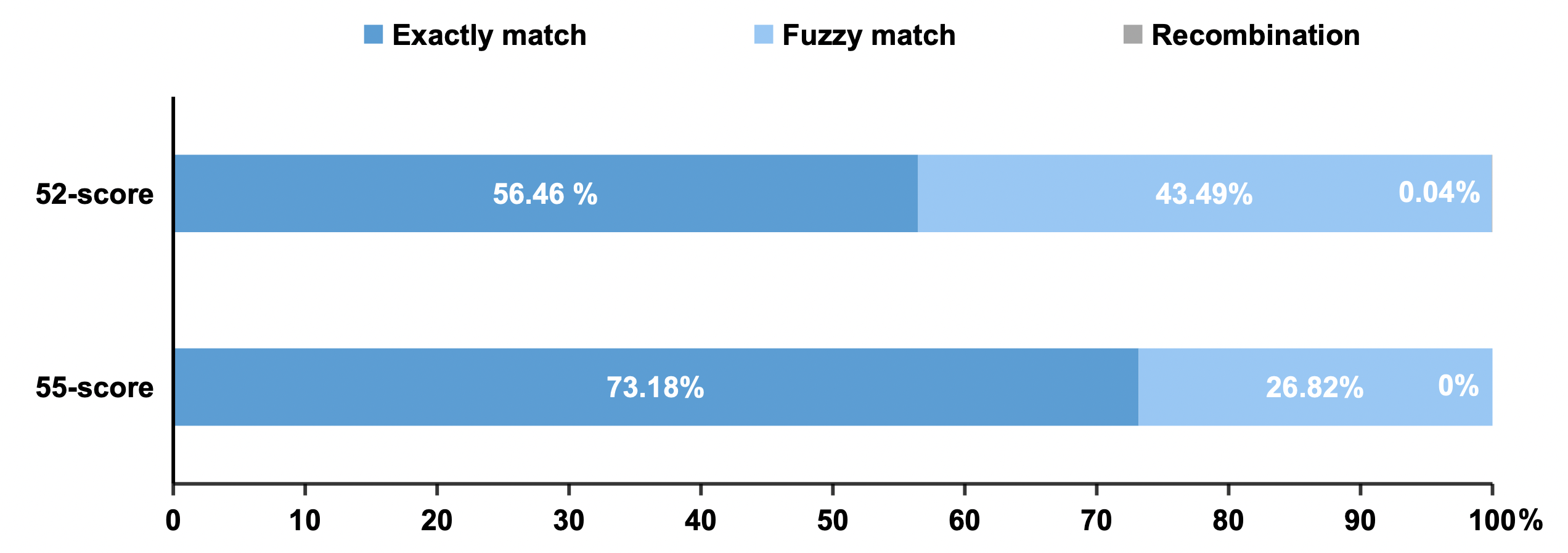

**Figure S4.** **Sequencing analysis of eight DNA-MT assembled blocks using linker combinations with scores of 55 and 52.**

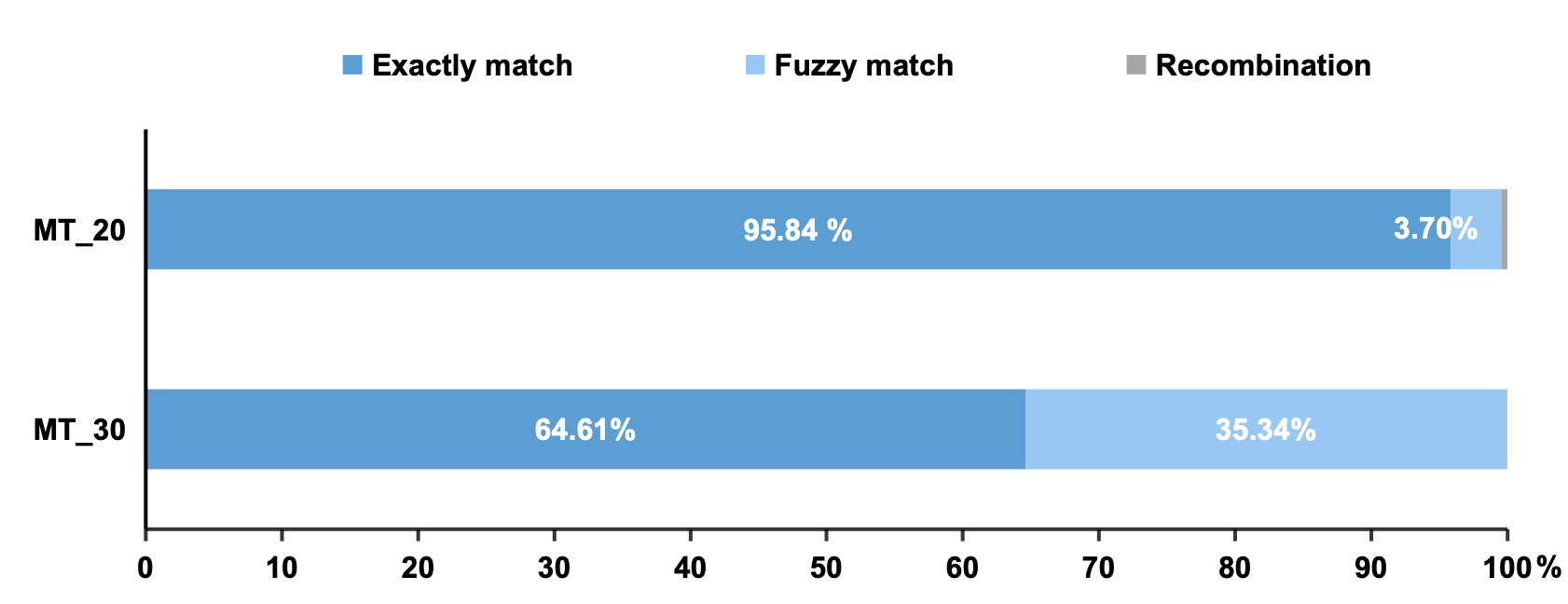

**Figure S5. Sequencing analysis of 12 DNA-MT blocks assembled using 20 bp and 30 bp information carriers.**

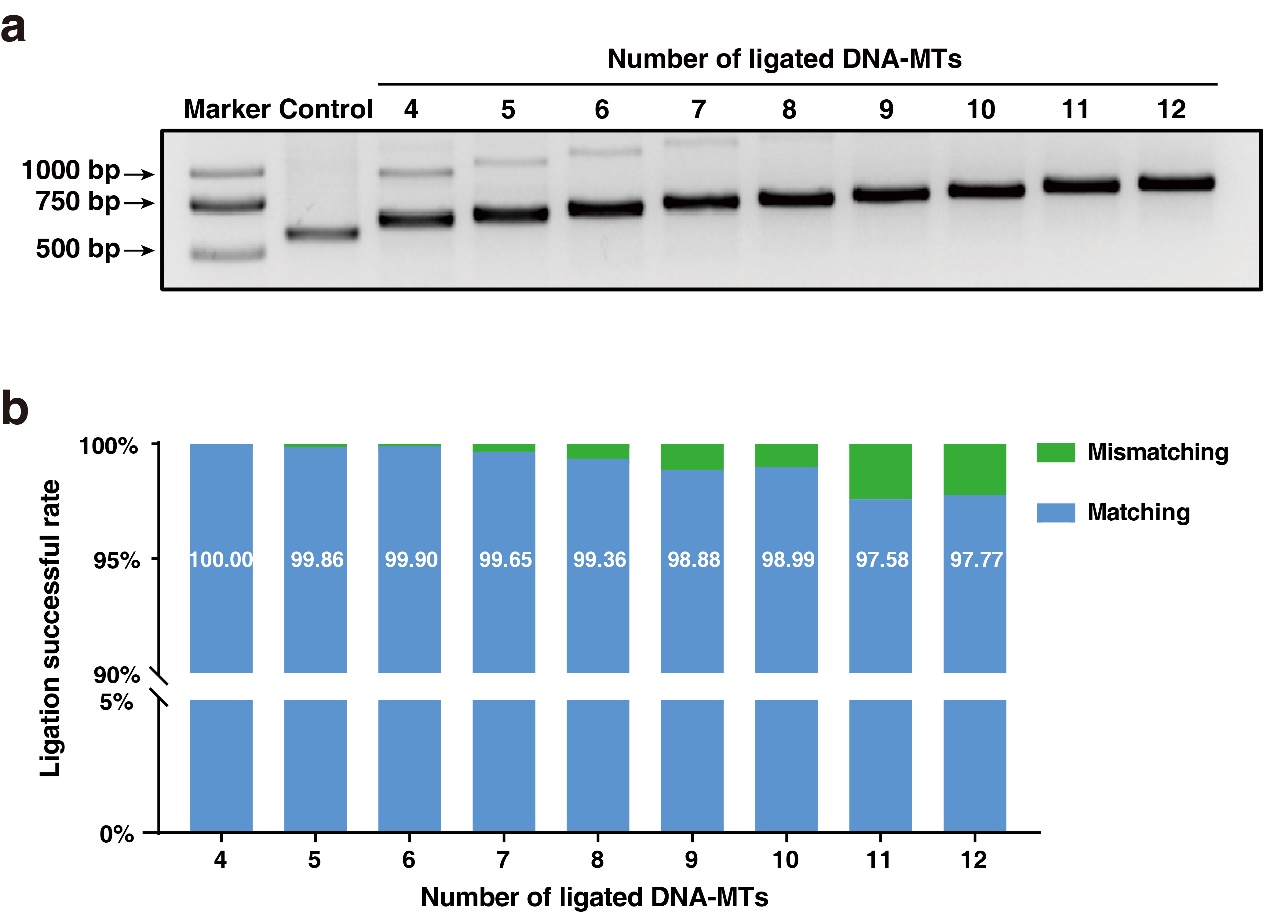

**Figure S6. Characterization of manually assembly of DNA movable type blocks using one-step enzymatic ligation. a, Agarose gel electrophoresis of DNA-MT block products from enzymatic ligation of 4 to 12 DNA-MTs.** Nine one-step ligation reactions were performed to assemble 4 to 12 DNA-MTs. The inserted DNA-MT blocks in recombinant plasmids were PCR (Test-F/Test-R) amplified, and the PCR products were run on a 1.5% agarose gel. Empty plasmids without any inserts served as controls, showing a 514 bp band. Inserts with 4 to 12 DNA movable types displayed distinct amplification bands at 561 bp, 585 bp, 609 bp, 633 bp, 657 bp, 681 bp, 705 bp, 729 bp, and 753 bp, respectively. Successful ligation was then confirmed by Sanger sequencing (Applied Biosystems 3730xl DNA Analyzer). **b, Sequencing of ligation products for assembled DNA-MT blocks using next-generation sequencing (Illumina NovaSeq6000).** Recombinant plasmids containing 4 to 12 DNA-MT inserts in E. coli were extracted for sequencing. The sequencing results showed that almost all sequences exactly matched the desired DNA-MT blocks, with only a very small number of sequences missing at least one DNA-MT.

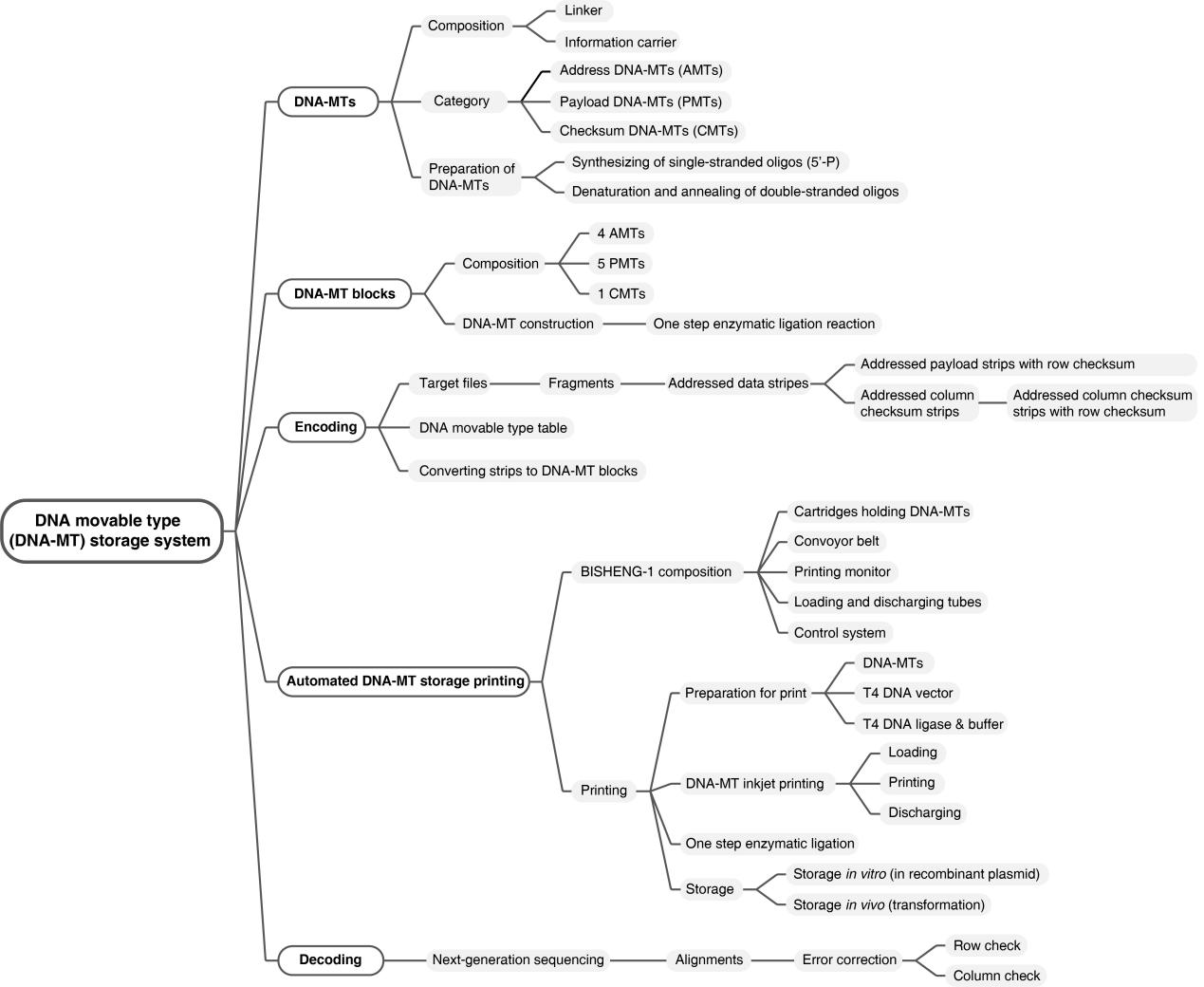

**Figure S7. Brain atlases of DNA movable type storage.** This map provides a comprehensive overview of the system, including the design of DNA-MTs, DNA-MT libraries (payload, address, and checksum), a coding-decoding system (comprising DNA-MT table and DNA-MT block assemblies), and an automatic DNA-MT inkjet printer, BISHENG-1

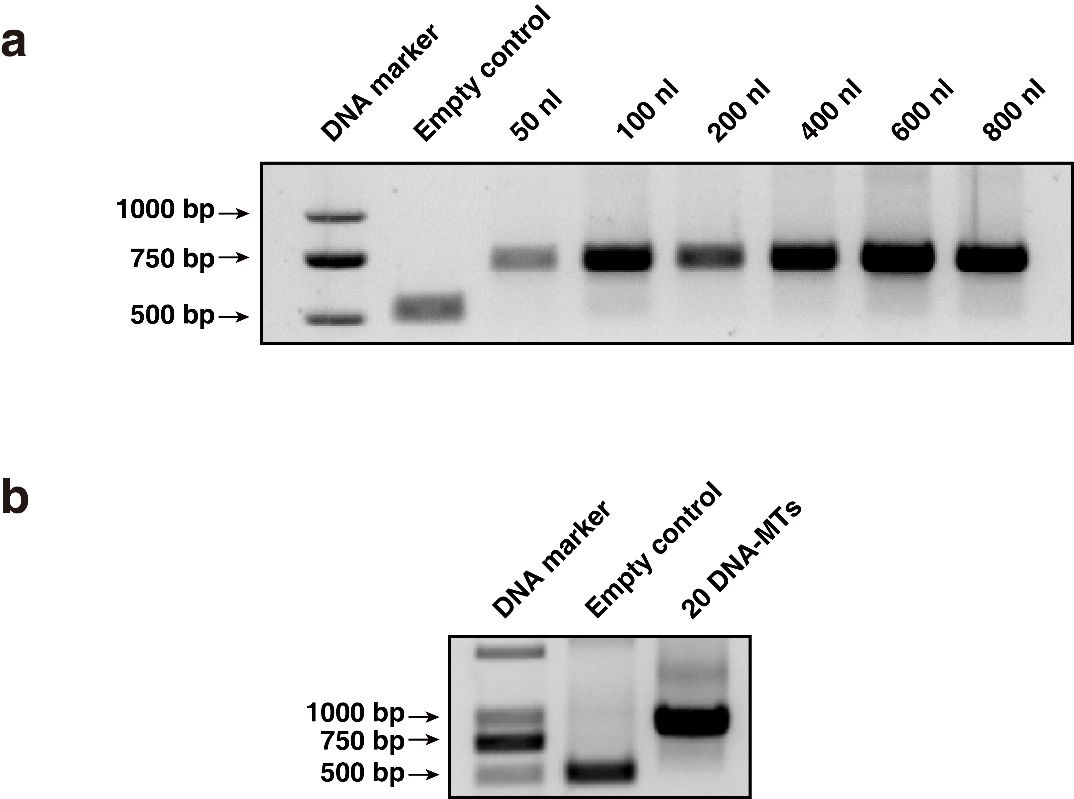

**Figure S8. Scalability testing of** **DNA-MT blocks assembly using one-step enzymatic ligation.** **a.** **Assembly of 10 DNA-MT blocks at nano-scale.** We employed acoustic droplet ejection (ADE) technology (ECHO650, Labcyte) to decrease the volume of DNA-MT droplets from 50 nl to 800 nl. The ligation products of 10 DNA-MT blocks assembled in recombinant plasmids were amplified via PCR using Test-F/Test-R primers and run on an 1% agarose gel. The PCR fragments measured 705 bp for the ligation product and 514 bp for the empty control. The accuracy of the assembly was subsequently verified by Sanger sequencing of the PCR-amplified fragments. **b, Assembly of 20 DNA-MT blocks.** The ligation product of 20 DNA-MTs (shown in Supplementary Table 3) was assembled in recombinant plasmids and initially transformed into E.coli for long-term preservation. These blocks were then PCR amplified using Test-F/Test-R primers and run on an 1% agarose gel. The PCR fragments measured 945 bp for the ligation product and 514 bp for the empty control. The assembly accuracy was then verified by Sanger sequencing of the PCR amplification fragments

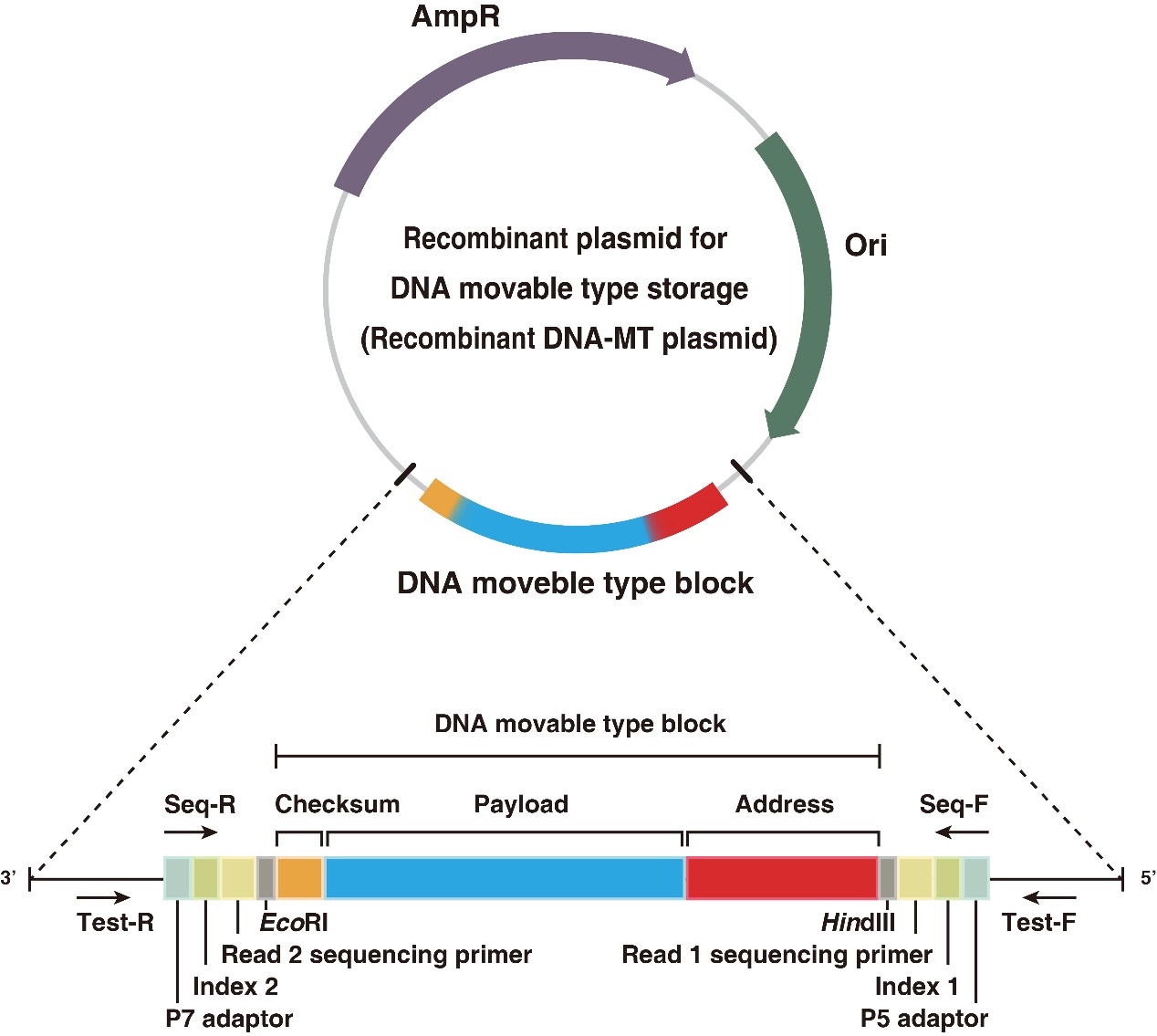

**Figure S9. Construction of DNA-MT plasmid.** This plasmid has been engineered to incorporate Illumina sequencing sequences to minimize sequencing costs (TableS3). Specifically, P5 adaptor, index 1, and Read 1 sequencing primer sequences were added adjacent to the 5’ end of the HindIII site, and P7 adaptor, index 2, and Read 2 sequencing primer sequences were added at the 3’ end of the EcoRI site. Amplification primers (Seq-F/Seq-R) were designed for direct library construction and sequencing of DNA-MT blocks on the Illumina platform

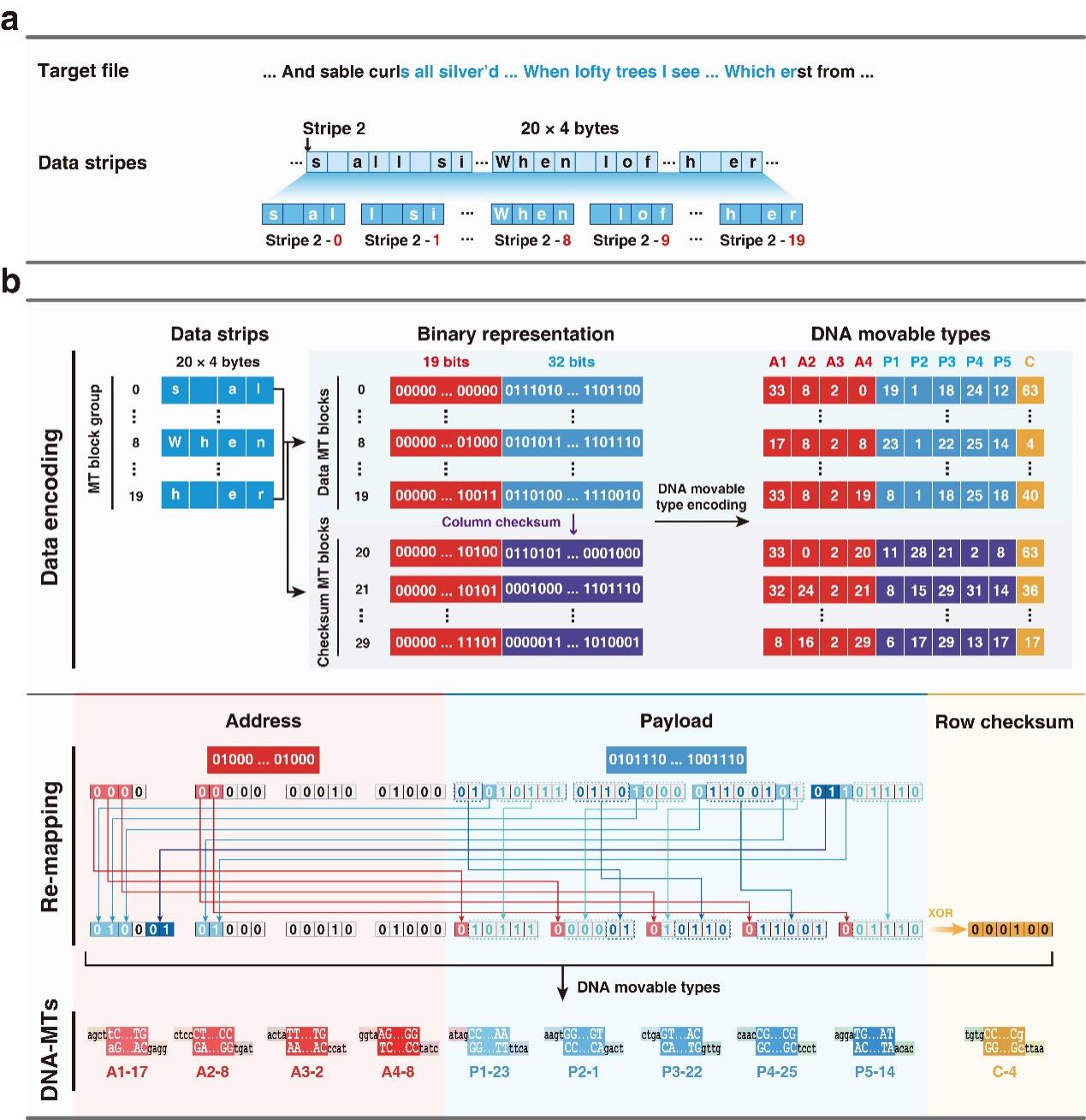

**Figure S10. Encoding the target file of Shakespeare's Sonnet 12 into DNA-MT blocks using BISHENG-1. a**, **Splitting the target file into data stripes.** We first split the target file into 80-character fragments and then divide each fragment into twenty 4-character data strips. **b, Encoding data strips into DNA movable type blocks.** For every 20 data strips, we generate 10 column-checksum strips are generated using the Reed-Solomon algorithm for error detection and correction. The original arrangement of BISHENG-1’s cartridges often result in the overuse of certain cartridges, thereby reducing their lifespan. To avoid this, we shuffle the cartridges by remapping the data and column-checksum strips. This remapping process includes exchanging the top 3 bits of the highest order AMT and the top 2 bits of the second highest order AMT with the top 1 bit of 5 PMTs. This strategy is designed to achieve a balanced use of cartridges across the library. A row-checksum is then calculated and appended to each strip before translating each strip into a DNA movable type block

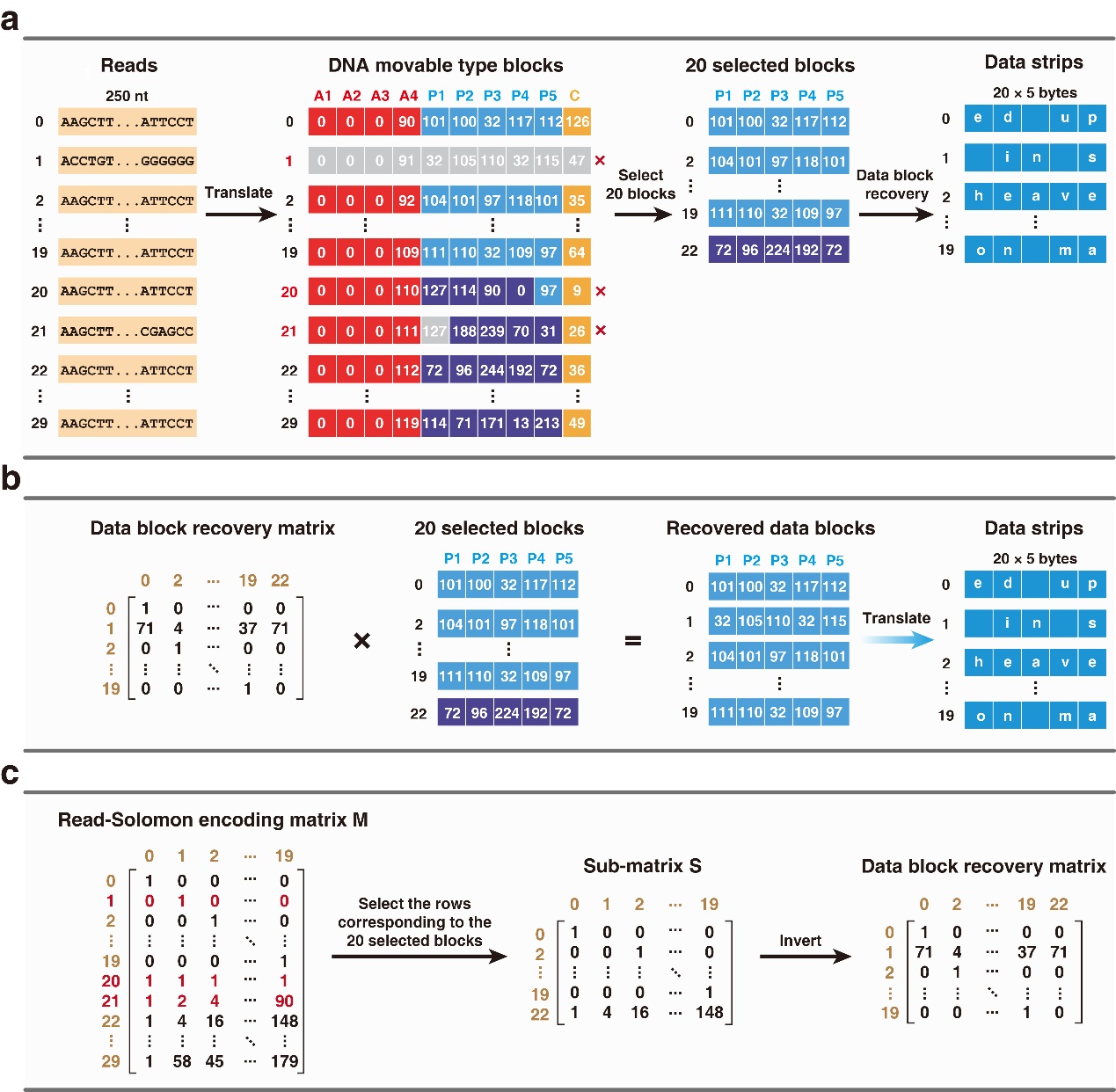

**Figure S11. Reconstructing original data from DNA movable type stored file. a, Decoding sequencing reads into data strips.** Sequencing reads are decoded into corresponding DNA movable types based on a DNA movable type code table (Supplementary Table 1), which are then compiled into data strips, further forming a data fragment (comprising 20 × 5 = 100 bytes). The missing data strips can be recovered using column-checksum DNA-MTs. In this case, the row 22 data strip, randomly selected, is used for recovery of row 1 data strip. **b, Recovering missing data blocks.** A set of 20 data blocks from a fragment (from 19 data strips and the row 22 column-checksum data strip) is selected and multiplied with a pre-calculated data block recovery matrix, translating recovered data blocks into the original 20 data strips (100 bytes). **c, Calculating the data block recovery matrix.** A 20×20 sub-matrix is extracted from a pre-calculated Reed-Solomon encoding matrix, and its inverse is calculated to serve as the data block recovery matrix
