## Supplementary figures and images for "Cost-effective DNA Storage System with DNA Movable Type"

File: 0001_32824041200720_(2.5P)_[881R].ab1 Run Ended: 2024-04-13 00:25:28 Signal G:484 A:446 T:425 C:491
Sample: 72801503391_32824041200720_(2.5P)_[881R]_PW Lane: 4 Base spacing: 16.06 679 bases in 8189 scans Page 1 of 1

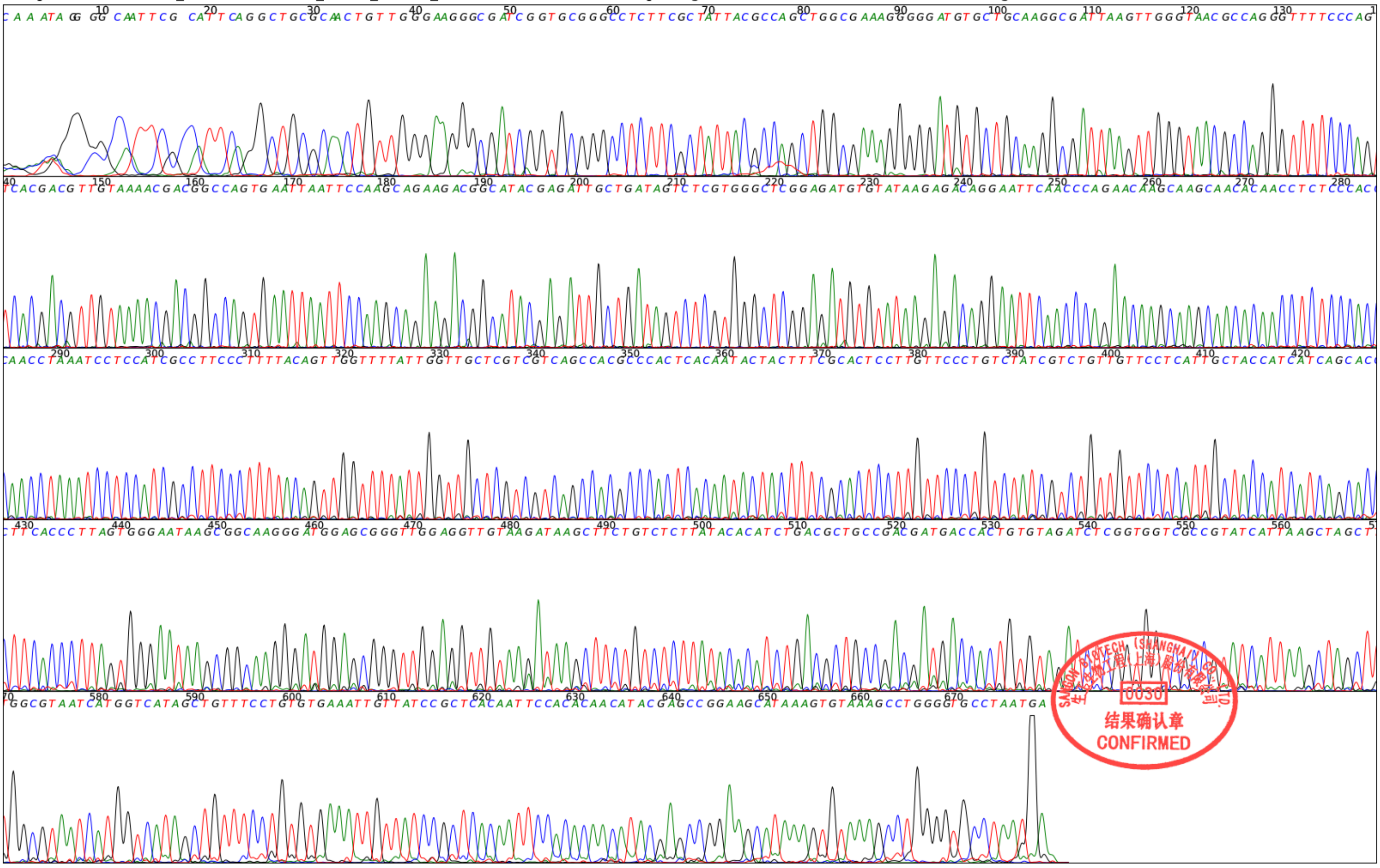

Supplement: Supplementary Data 2 [file 603163_file08.zip › 0001_32824041200720_(2.5P)_[881R].pdf]

File: 0002_32824041200721_(5P)_[881R].ab1 Run Ended: 2024-04-13 00:25:28 Signal G:1027 A:956 T:979 C:1153

Sample: 72801503392_32824041200721_(5P)_[881R]_PW Lane: 4 Base spacing: 16.05 678 bases in 8166 scans 

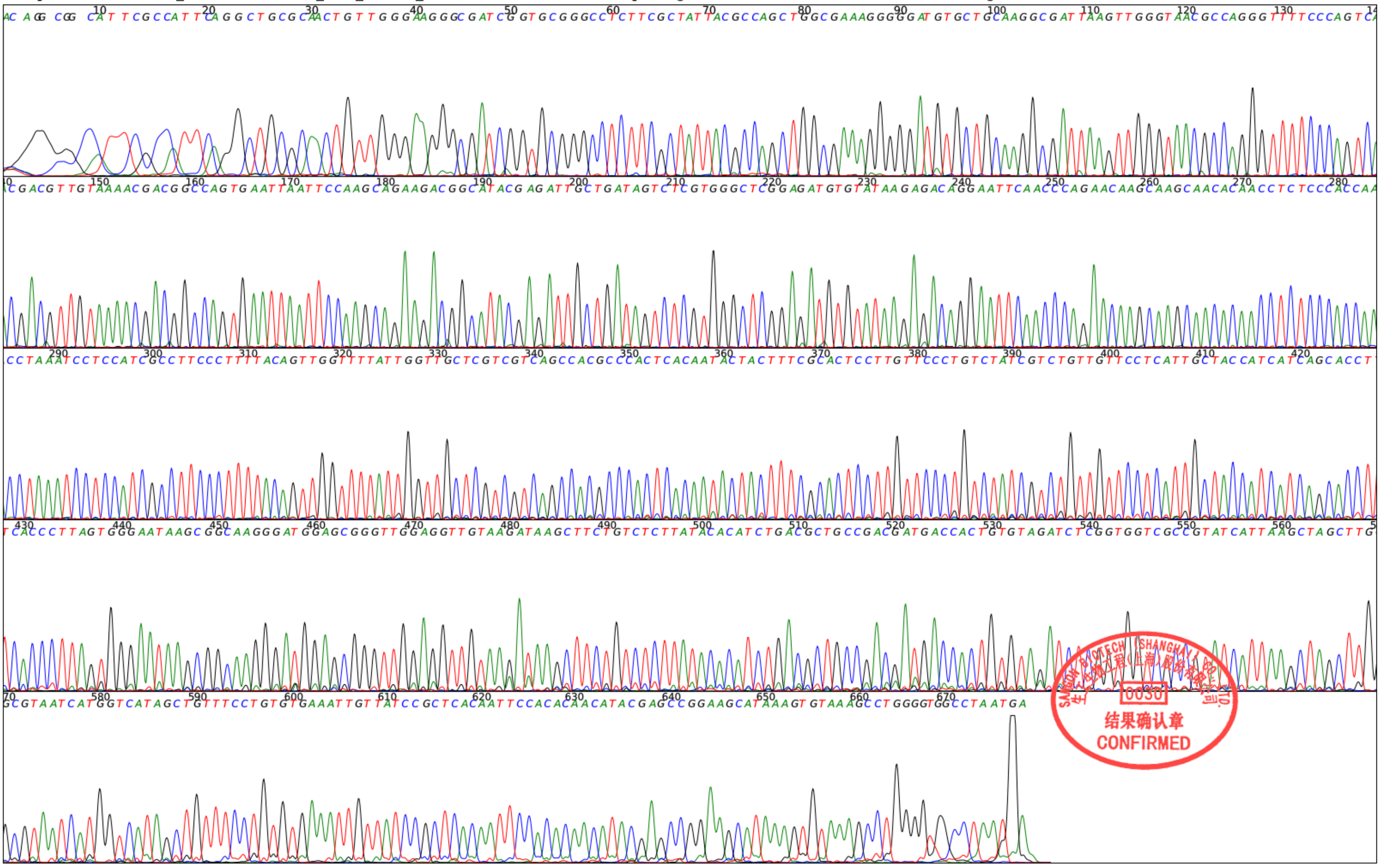

Supplement: Supplementary Data 2 [file 603163_file08.zip › 0002_32824041200721_(5P)_[881R].pdf]

File: 0003_32824041200722_(10P)_[881R].ab1 Run Ended: 2024-04-13 00:25:28 Signal G:2507 A:2779 T:2787 C:3103

Sample: 72801503393_32824041200722_(10P)_[881R]_PW Lane: 4 Base spacing: 15.98 671 bases in 8182 scans Page 1 of 1

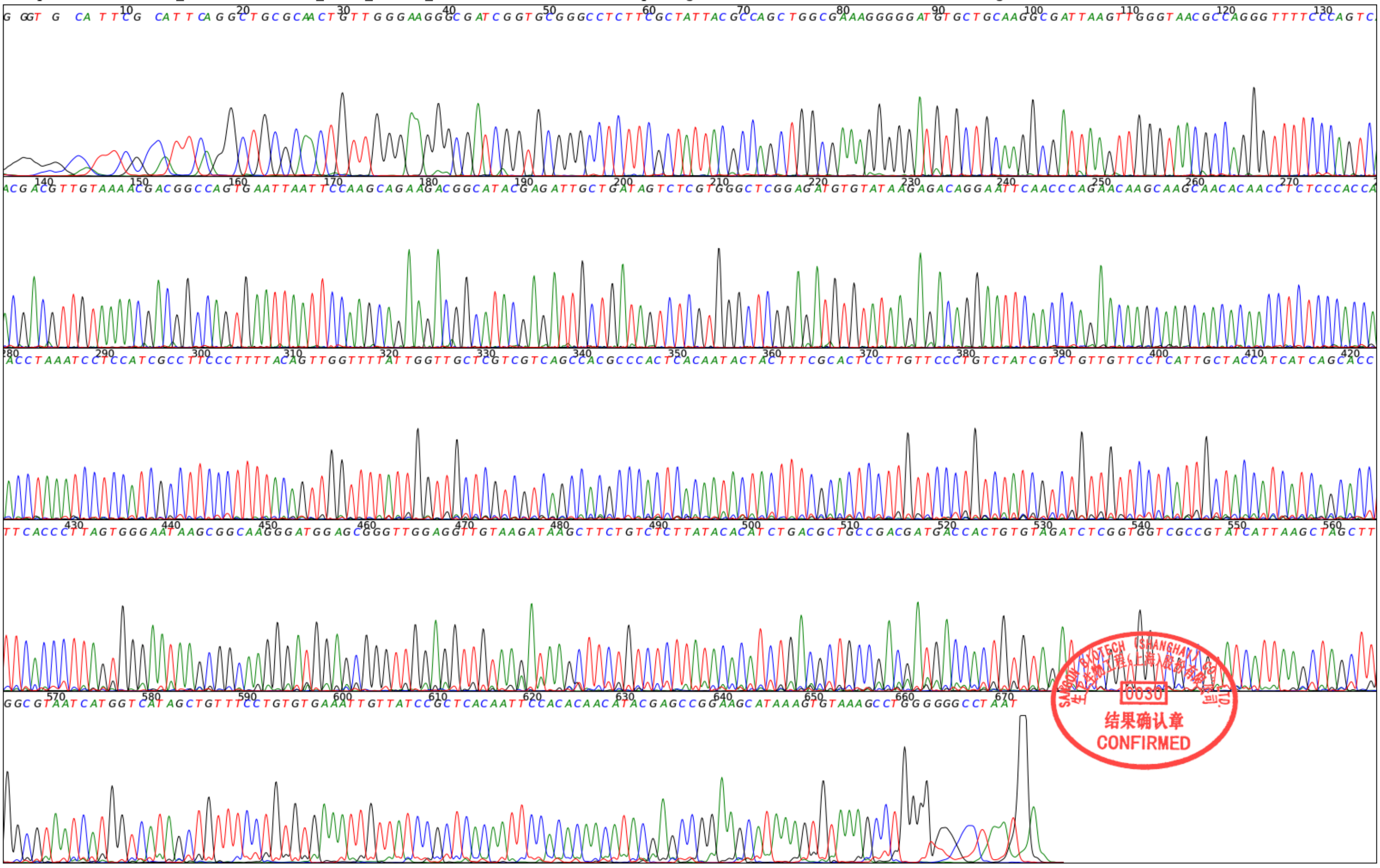

Supplement: Supplementary Data 2 [file 603163_file08.zip › 0003_32824041200722_(10P)_[881R].pdf]

File: 0004_32824041200723_(20P)_[881R].ab1 Run Ended: 2024-04-13 00:25:28 Signal G:1808 A:2046 T:2250 C:2574

Sample: 72801503394_32824041200723_(20P)_[881R]_PW Lane: 4 Base spacing: 15.99 675 bases in 8197 scans Page 1 of 1

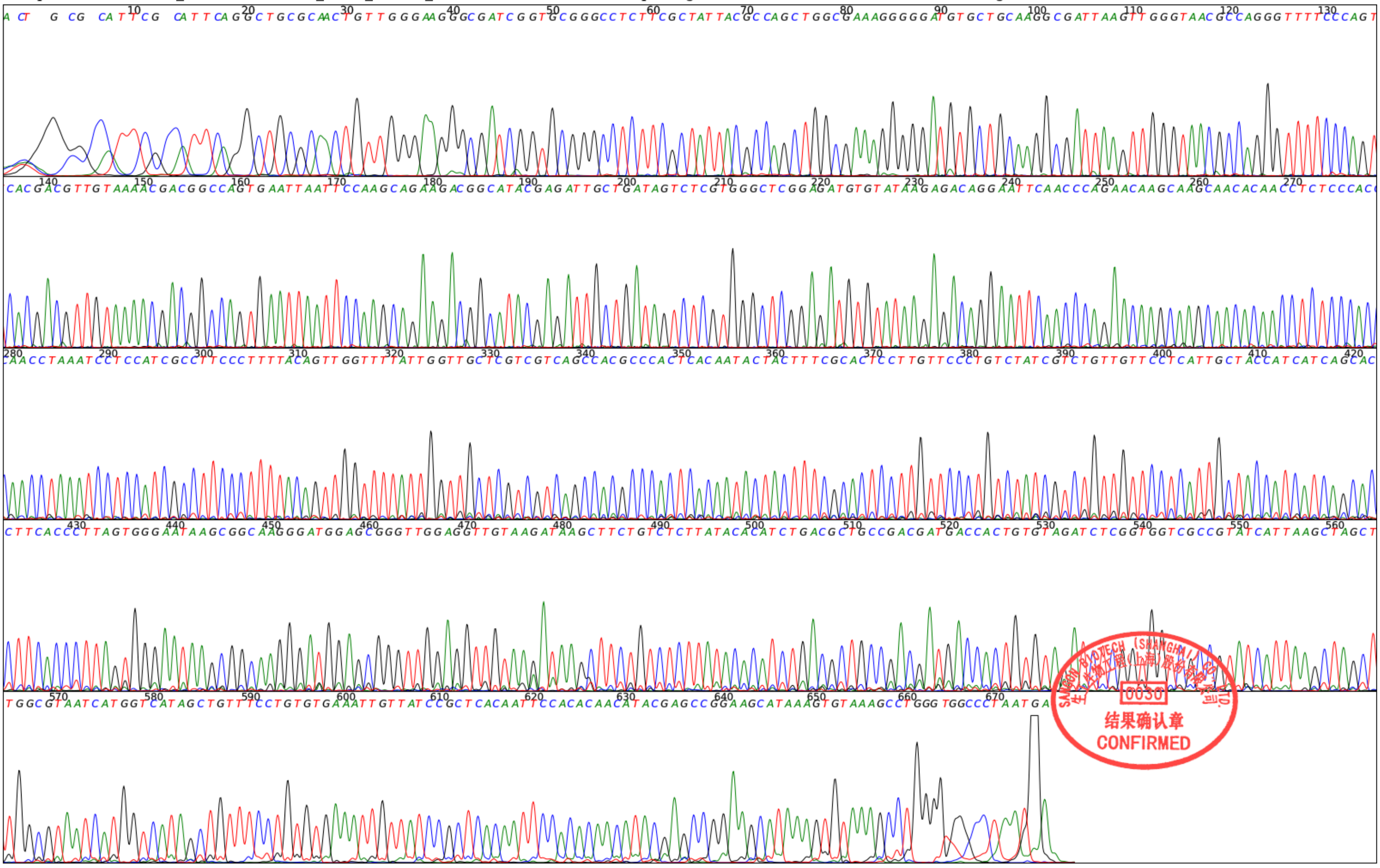

Supplement: Supplementary Data 2 [file 603163_file08.zip › 0004_32824041200723_(20P)_[881R].pdf]
